# Structural basis for catalysis and selectivity of phospholipid synthesis by eukaryotic choline-phosphotransferase

**DOI:** 10.1101/2024.05.09.593427

**Authors:** Jacquelyn R. Roberts, Yasuhiro Horibata, Frank E. Kwarcinski, Vinson Lam, Ashleigh M. Raczkowski, Akane Hubbard, Betsy White, Hiroyuki Sugimoto, Gregory G. Tall, Melanie D. Ohi, Shoji Maeda

**Author notes:** J.R.R., Y.H., F.E.K. contributed equally. J.R.R. and Y.H. have the right to list oneself first in bibliographic documents. M.D.O. and S.M. are joint corresponding authors. Xaira Therapeutics, Brisbane, CA, 94005, USA.

## Abstract

Phospholipids are the most abundant component in lipid membranes and are essential for the structural and functional integrity of the cell. In eukaryotic cells, phospholipids are primarily synthesized *de novo* through the Kennedy pathway that involves multiple enzymatic processes. The terminal reaction is mediated by a group of cytidine-5’-diphosphate (CDP)-choline /CDP-ethanolamine-phosphotransferases (CPT/EPT) that use 1,2-diacylglycerol (DAG) and CDP-choline or CDP-ethanolamine to produce phosphatidylcholine (PC) or phosphatidylethanolamine (PE) those are the main phospholipids in eukaryotic cells. Here we present the structure of the yeast CPT1 in multiple substrate-bound states. Structural and functional analysis of these binding-sites reveal the critical residues for the DAG acyl-chain preference and the choline/ethanolamine selectivity. Additionally, we present the structure in complex with a potent inhibitor characterized in this study. The ensemble of structures allows us to propose the reaction mechanism for phospholipid biosynthesis by the family of CDP-alcohol phosphatidyltransferases (CDP-APs).

---

Cellular membranes, made of lipid bilayers, create a physical barrier separating the intracellular and extracellular environments and encapsulate subcellular organelles to create intracellular compartments^1^. Membranes serve as a platform where numerous physiological processes are regulated via integral- or peripheral-membrane proteins^2^. The lipid bilayer is composed primarily of phospholipid molecules. Phosphatidylethanolamine (PE) is the main phospholipid in prokaryotic cells and phosphatidylcholine (PC) is the main phospholipid in eukaryotic cells. PC accounts for up to 50% of total eukaryotic cellular phospholipids^1,3^ Besides their role as a major structural component of cell membranes, phospholipids serve as precursor substrates for signaling molecules such as lysophosphatidic acid or platelet-activating factor^4^.

Given the fundamental roles phospholipids have in numerous cellular processes, coordinated synthesis and metabolism of these molecules are essential for normal cellular physiology and the regulation of numerous biological processes^5^. PE and PC are synthesized *de novo* through a three-step enzymatic process known as the Kennedy pathway^6,7^. In the first step, ethanolamine or choline is phosphorylated to form phosphoethanolamine or phosphocholine. In the second step, the phosphorylated molecules are coupled with cytidine-5’-triphosphate (CTP) to form cytidine-5’-diphosphate (CDP)-ethanolamine or CDP-choline, high-energy intermediates. In the third step, CDP-ethanolamine or CDP-choline are conjugated with diacyl-glycerol (DAG) to form PE or PC (Supplementary Fig. 1). In mammalian cells, the majority of PE and PC are synthesized through the Kennedy pathway. This enzymatic pathway is ubiquitously present and conserved in eukaryotes, including *Saccharomyces cerevisiae*, making yeast a powerful model system for investigating lipid metabolism^8^.

Members of the integral membrane CDP-alcohol phosphotransferase (CDP-AP) family mediate the final enzymatic step of the Kennedy pathway. Ethanolaminephosphotransferase (hCEPT1 and hEPT1 in human and yEPT1 in yeast) conjugates CDP-ethanolamine to DAG to form PE, while Cholinephosphotransferase (hCHPT1 in human and yCPT1 in yeast) conjugates CDP-choline with DAG to form PC^9–12^ (Supplementary Fig. 1). CDP-APs have a universally conserved signature motif, (D(x)_2_DG(x)_2_AR(x)_7,8_G(x)_3_D(x)_3_D), that plays a critical role in catalyzing the transfer of a chemical moiety with a phosphate group from a CDP-linked donor to an alcohol acceptor in a divalent cation-dependent manner. Mutation of any residues in the signature motif reduces the catalytic activity of the protein^13^. Cholinephosphotransferases and ethanolaminephosphotransferases are highly conserved with hCHPT1 and hCEPT1 sharing 62% sequence identity and yeast yCPT1 and yEPT1 sharing 55% sequence identity. However, these enzymes have different substrate specificities. hCHPT1 and yCPT1 have specificity for CDP-choline, while hCEPT1 and yEPT1 are more promiscuous enzymes capable of catalyzing both CDP-choline and CDP-ethanolamine^14–16^. In addition to the selectivity for the lipid headgroup, yCPT1 and yEPT1 also show different preferences toward DAG acyl-chain group species^14^.

Several structures of prokaryotic CDP-APs have been determined^17–23^. They all form dimers with the dimerization interface at the transmembrane (TM) helices 3 and 4. These structures provided valuable insights into the overall architecture, molecular basis of substrate recognition, and the catalytic mechanism of this family of integral membrane enzymes. Recently, two structures of eukaryotic CDP-APs, human CEPT1 and *Xenopus laevis* CHPT1, were determined^24,25^. The structures showed the overall architecture of each enzyme, the binding mode of PC, and suggested possible substrate entry sites. They also highlighted that while eukaryotic and prokaryotic CDP-APs shared many conserved structural features, including being dimers and a similar mode of CDP binding, there were also key differences, such as having different dimer interfaces. However, none of the structures provided molecular insight into how enzymes that share such a high degree of sequence similarity and structural conservation have different substrate selectivity.

Here we present the structures of full-length yCPT1 in different substrate-bound states determined by single particle cryo-electron microscopy (cryo-EM). These structures show that, like other CDP-APs, yCPT1 forms a dimer although using an entirely different interface than seen in the known prokaryotic or eukaryotic structures^17–25^. Our structures of yCPT1 in complex with either DAG or CDP-choline delineate the binding mode of these substrates and mutational analyses identified the residues critical for the DAG acyl-chain preference and head-group selectivity. We also found that chelerythrine, previously reported as a hCEPT1 inhibitor^26^, potently inhibits yCPT1. The cryo-EM structure of yCPT1 bound with chelerythrine reveals the inhibitory mechanism of this molecule and the comparison with the structure of yCPT1 bound to CDP-choline provides molecular insights into the catalytic mechanism of this enzyme.

## Structural characterization of yCPT1 and comparison with other CDP-AP structures

*S. cerevisiae* yCPT1 was expressed and purified from Sf9 insect cells (Supplementary Fig. 2a). The purified protein is enzymatically active as determined by thin-layer chromatography (TLC) using radiolabeled CDP-choline (Supplementary Fig. 2b). Unexpectedly, the addition of CDP-choline without adding DAG still produced PC, indicating that purified yCPT1 contains co-purified endogenous DAG (Supplementary Fig. 2c). Using single-particle cryo-EM analysis, we determined a 3.2 Å resolution structure of yCPT1 bound to a nanobody (Nb24) isolated from a synthetic nanobody library^27^ (Fig. 1a, Supplemental Fig. 2d and 3, and Supplementary Table1). Nb24 binding did not alter yCPT1 enzymatic activity (Supplemental Fig. 2e) and the diffuse density for the Nb24 visible in the 2D averages suggests that this nanobody interacts with a flexible lumenal loop of yCPT1 (Supplemental Fig. 3b-d). The quality of the density map allowed us to build an atomic model of the yCPT1 dimer (Fig. 1b); however, the fragmented density corresponding to the nanobody kept us from modeling this part of the structure (Supplementary Fig. 3g).

**Fig. 1.**
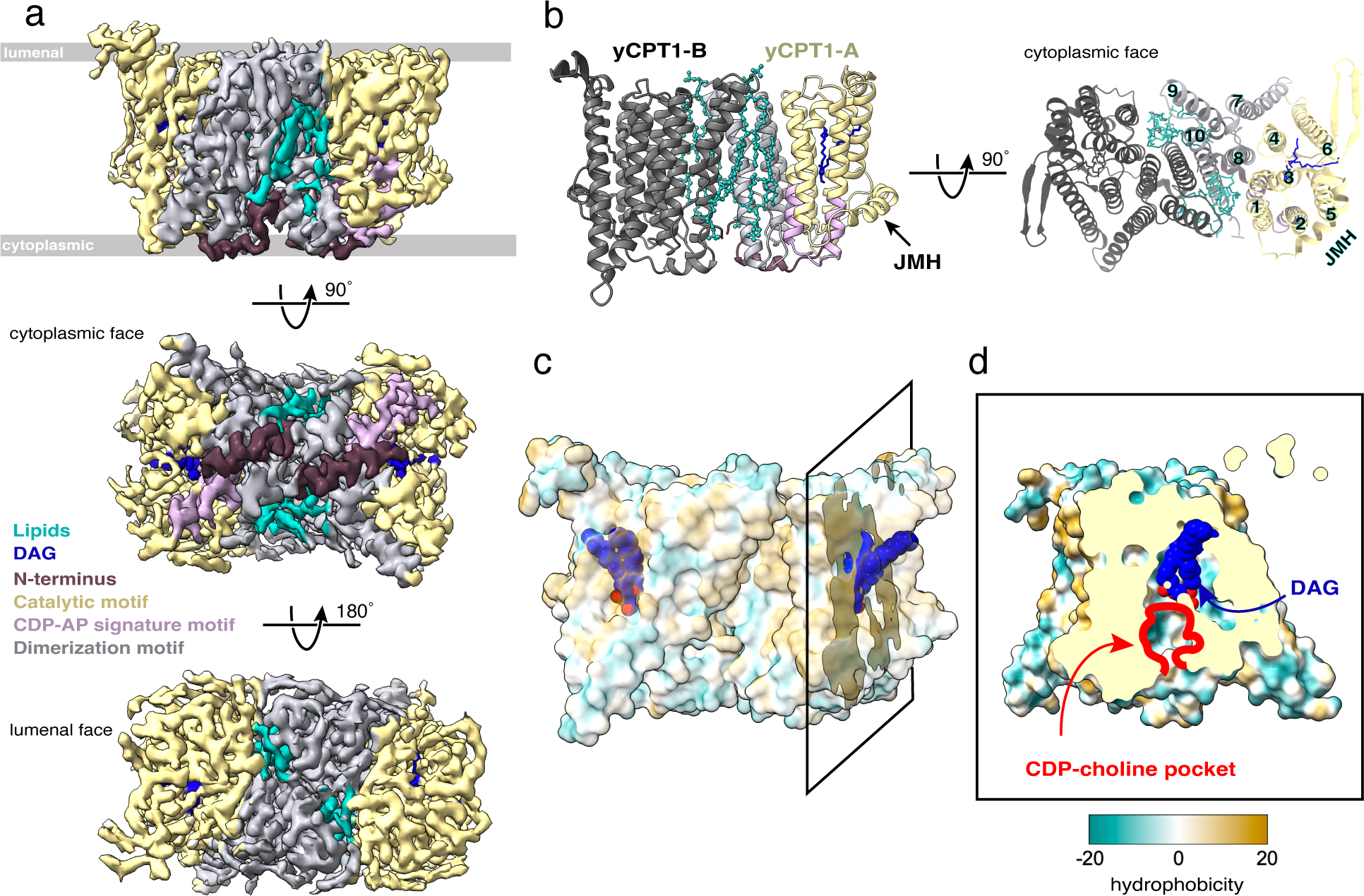
Overall architecture of yCPT1. **a.** 3.2 Å resolution cryo-EM density map of dimeric yCPT1. Each ligand, motif, and assembly are colored according to the panel legend: yellow, catalytic assembly; purple, CDP-AP signature motif; grey, dimerization assembly; brown, N-terminal stretch; cyan, phosphatidylcholine (PC); blue, diacylglycerol (DAG). Positions of lumenal and cytoplasmic membranes shown in top panel. **b.** Model of dimeric yCPT1. Atomic models corresponding to the density in panel **a**. In one protomer of the model each ligand, motif, and assembly are colored similarly as in panel **a**. In the lumenal view, the TM helix is numbered and the juxtamembrane helix (JMH) is labeled. Inter-monomer PCs and DAG are depicted as sticks. **c.** Surface model of yCPT1 colored with hydrophobicity index by Chimera^50^. DAG is shown as balls within the surface model colored in blue(carbon) and red(oxygen). The cross-section plane depicted in panel d is also shown. **d.** Cross-section of yCPT1 at the substrate-binding pocket. CDP-choline-binding site is enclosed in a solid red line. The color code of the models and the hydrophobicity index of yCPT1 is the same as shown in the panel c.

The yCPT1 dimer is composed of two protomers of 10 TM helices with both N- and C-terminus located intracellularly. The juxtamembrane helix (JMH) precedes the TM helices and is located at the peripheral edge of the protein (Fig. 1b, Supplementary Fig. 4a). The catalytic motif is composed of TM helices 1-6 and contains the substrate binding pocket and potential substrate entry site. At its catalytic center, yCPT1 possesses a conserved CDP-AP signature motif (^96^**<u>D</u>**AC**<u>DG</u>**MH**<u>AR</u>**RTGQQGPL**<u>G</u>**ELF**<u>D</u>**HCI**<u>D</u>**^121^) that spans from the lower half of TM helix 2 to the middle of TM helix 3 (Fig. 1a, b, Supplementary Fig. 4a). Metal ions, such as Ca^2+^, Zn^2+^, Mn^2+^, and Na^+^, are coordinated with the four aspartate residues in the signature motif of various prokaryotic CDP-APs. Because yCPT1, like xlCHPT1 and hCHPT1, requires Mg^2+^ or Mn^2+^ for activity^25,28,29^ and we included 10mM MgCl_2_ in the sample buffer, we modeled Mg^2+^ into the densities observed at the catalytic center of each protomer (Supplementary Fig. 4b).

As seen with other eukaryotic CDP-APs^24,25^, the TM helices that form the catalytic motif of yCPT1 superimpose onto the six TM helices found in prokaryotic CDP-APs (*e.g.*, r.m.s.d. = 2.85 Å between yCPT1 and *af*DIPPS, a representative prokaryotic CDP-AP from archaea, PDBID: 4MND) (Supplementary Fig. 4d,e). Eukaryotic CDP-APs have an extra four helical bundle, TM helices 7-10, in addition to the six TM helices that form the catalytic domain (helices 1-6, Fig. 1b and Supplementary Fig. 4a, d-f). These additional TM helices contribute to the dimerization motif of yCPT1 (Fig. 1a,b). The yCPT1 dimer is composed of a large interface formed by extensive protein-protein and lipid-protein interactions. yCPT1 TM helices 9 and 10, with a smaller contribution from TM helix 1, support protein interactions between protomers that span from the lumenal to the cytoplasmic face of the dimer (Fig. 1, Supplementary Fig. 4a, d). Additionally, three molecules of phospholipid density were observed filling a groove formed at the interface of the monomers and appear to stabilize the interactions between these protomers (Supplementary Fig 4c). To examine the dimer interface more closely, we calculated the inter-subunit surface areas occupied by either lipid- or protein-mediated contacts using the PDBePISA server^30^. This analysis shows that the lipid-mediated contact area is ∼1100 Å^2^, while the protein-protein contact surface area is ∼940Å^2^. These calculations indicate that lipid-protein and protein-protein contacts have equally important roles in stabilizing yCPT1 dimerization (Supplementary Fig. 5)^15^. Interestingly, previous studies reported that the supplementation of phospholipids to solubilized yCPT1, particularly PC, increases its activity in *in vitro* assays^15^

While the CDP-AP family appear to be obligate dimers, structural analyses show that the dimerization interfaces are surprisingly not conserved among eukaryotes or between eukaryotes and prokaryotes. For all reported prokaryotic CDP-AP structures, the dimerization interface involves interactions between TM helices 3 and 4 of each protomer. The structures of the eukaryotic hCEPT1 and xlCHPT1 enzymes have a dimerization interface mediated by a small area of interaction at the lumenal side between TM helices 7 and 9^24,25^. The hCEPT1 and xlCHPT1 dimerization motif is smaller than that seen in yCPT1 and composed mostly of protein-protein contacts, although there were also inter-dimer phospholipid molecules observed at the dimer interface in the xlCHPT1 structure^25^. Thus, these analyses show that while CDP-APs are structurally conserved and form dimers, the dimerization interface and the contributions of lipids to support dimerization are not conserved between species (Supplementary Fig. 4d-f).

## DAG binding site and acyl-chain selectivity

In addition to the densities for the lipids found at the dimerization interface, the cryo-EM density map of yCPT1 also revealed an elongated, V-shaped density in the hydrophobic channel near the catalytic center of each protomer. We assigned this density as DAG (Figs. 1c,d and 2a-c). Evidence supporting this modeling includes the observation that purified yCPT1 produces PC upon addition of CDP-choline without adding extra DAG (Supplementary Fig. 2c) combined with mass spectrometry analysis of purified yCPT1 that detected DAGs with different acyl-chain length such as 16:1-16:1 and 18:1-18:1 (Supplementary Fig. 6c) and phospholipids including PC and PE with varying acyl-chain species ranging from 32:2 to 36:2 (Supplementary Fig. 6a,b). While the density could accommodate either a C16- or C18-DAG species, we modeled 16:1-16:1-DAG because a previous report indicated that yCPT1 prefers C16-DAG as a substrate rather than longer acyl-chain DAGs preferred by yEPT1^14,16^. In this model the sn-2 acyl-chain of the C16-DAG extends from around the N-terminal end of TM helix 5, passing through a channel between TM helices 5 and 6, and egresses towards the lipid environment (channel 2) (Fig. 2b-d). The sn-1 acyl-chain of the C16-DAG runs along TM helices 3 and 4 and exits between TM helices 4 and 6 (channel 1); however, unlike channel 2 that is exposed to the lipid environment, the exit of channel 1 is capped by a bulky phenylalanine side chain (Phe146, Fig. 2b-d). While the amino acids around residue 146 of yCPT1 and yEPT1 have high sequence conservation, residue 146 is not conserved. Specifically, yEPT1 has a less bulky leucine at this position (Leu146, Supplementary Fig. 7a), suggesting a structural explanation for the different DAG acyl-chain length specificities of yCPT1 and yEPT1. To examine the contribution of this residue in substrate specificity for these enzymes, we generated a predicted yEPT1 model using AlphaFold 2 (AF2)^31^ and compared it to the model built from the single particle cryo-EM yCPT1 map. This analysis shows that having a leucine at position 146 in yEPT1, rather than the Phe146 found in yCPT1, would open channel 1 to the membrane environment (Supplementary Fig. 7b). Comparison of the structures suggests a model where the bulkiness of the side-chain at the channel 1 exit directly influences acyl-chain length that can be accommodated via steric interference. To test this hypothesis, we replaced yCPT1 Phe146 with leucine and then evaluated the acyl-chain preference of this mutant enzyme (yCPT1 F146L) compared to wild type (WT) yCPT1. This analysis showed that as expected WT yCPT1 prefers C16-DAG as a substrate^14,16^, but yCPT1 F146L prefers C18-DAG (Fig. 2e). Additionally, mass spectrometry analysis of purified yCPT1 F146L detected an increased abundance of long-chain acyl DAGs compared to WT yCPT1 while PC and PE contents were unchanged (Supplementary Fig. 6d-f). Collectively, these analyses support a mechanism in which the bulkiness of the residue in this position of the enzyme (residue 146 in yCPT1 and yEPT1) sterically influences the acyl-chain length that can fit into channel 1. This finding explains the differences in DAG acyl length binding preferences of yCPT1 and yEPT1.

**Fig. 2.**
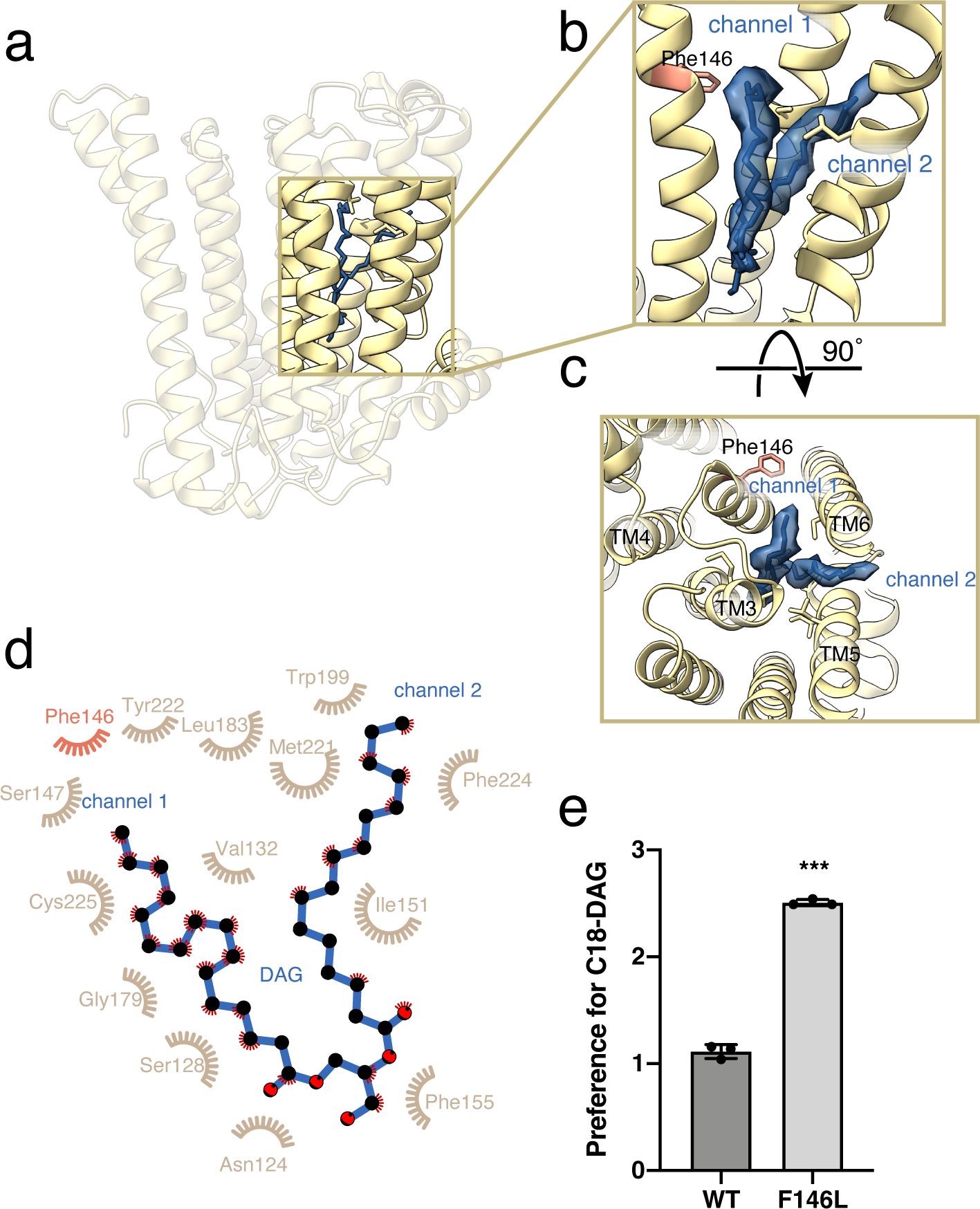
DAG binding mode of yCPT1. **a.** Model of yCPT1 protomer as viewed from the membrane plane. The box highlights the position of the DAG-binding site and is shown expanded in panels b and c. The cryo-EM density of DAG is shown in blue with the model also shown in blue. Acyl-chains from DAG occupy the hydrophobic spaces inside of yCPT1. **b,** Enlarged view of DAG-binding site. Each of the two acyl-chains of DAG occupies two distinct hydrophobic channels: Channel 1, the channel between TM4 and TM6; Channel 2, the channel between TM5 and TM6. **c.** The same DAG-binding site is viewed from the lumenal side. Note that Phe146 shown as sticks is located at the exit site of channel 1. **d.** DAG-surrounding residues in yCPT1. LigPlot2^51^ is used for the analysis and the illustration. Phe146 is shown in red. **e.** Choline phosphotransferase activity of yCPT1 WT and yCPT1 F146L mutant was measured using radiolabeled CDP-choline and DAG 16:0-16:0 or DAG 18:1-18:1 as the substrates. The preference for C18-DAG was determined by dividing the amount of PC 18:1-18:1 by PC 16:0-16:0. Data are presented as the mean ± SD. *** indicates P < 0.001 as compared with WT yCPT1. Statistical significance was determined using a Student’s t-test.

## CDP-choline binding site and substrate selectivity

To further understand the molecular basis of substrate recognition of eukaryotic CDP-APs enzymes, we next determined a structure of yCPT1 in the presence of 1 mM CDP-choline (Supplementary Fig. 8 and Supplementary Table1). The resulting 2.8 Å resolution map has clearly resolved density for both CDP-choline and PC, the end-product of the reaction, localized in the same binding pocket where DAG was found in the yCPT1 map determined in the absence of added CDP-choline (Fig. 3a-c). When comparing the positions of CDP-choline and PC, the β-phosphate moves ∼8 Å towards TM helix 4 with the choline group pointing toward the cytoplasmic side instead of pointing towards JMH (Fig. 3b,d). In a typical enzymatic cycle, the PC end-product is released from the catalytic site allowing a new DAG molecule to be bound. It is not clear from this structure why the PC molecule remains in the pocket and is not released. One explanation may be that the CDP-choline in the catalytic site stabilizes the β-sheet near the DAG binding pocket (Fig. 3b,d). In support of this model, the local B-factor distribution shows that in the absence of CDP-choline this area of yCPT1 is one of the most flexible regions of the structure, while the local B-factor distribution in the presence of CDP-choline shows that this part of the structure has decreased flexibility (Supplementary Fig. 9a,b). The stabilized β-sheet, which forms part of the cleft postulated to be a DAG entry site^25^, might make the exchange of DAG for PC more difficult *in vitro* than what happens in a physiological environment.

**Fig. 3.**
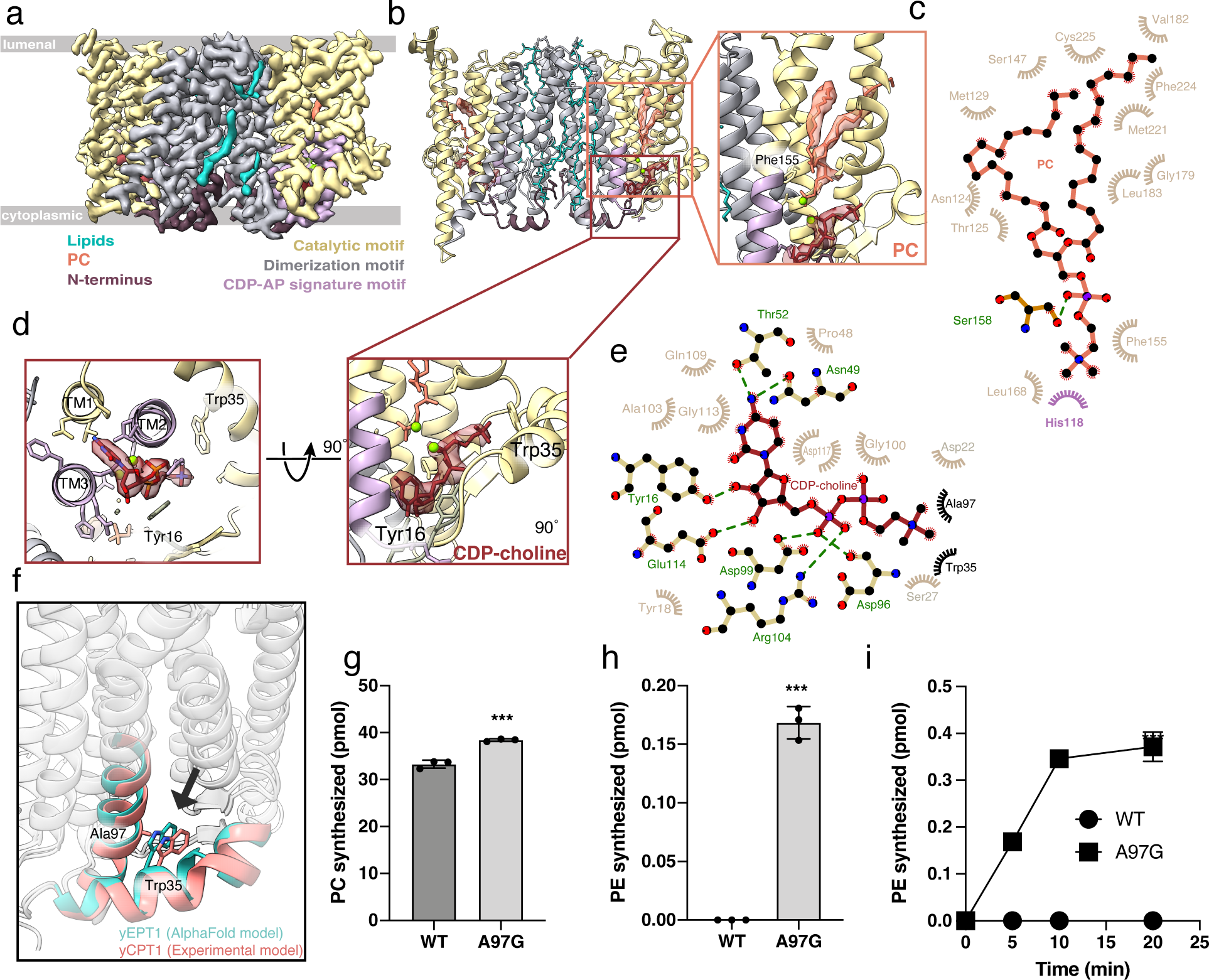
CDP-choline and PC binding mode. **a, b.** 2.9 Å resolution cryo-EM density map of yCPT1 dimer bound with CDP-choline and PC. Density is colored in the same way as in Fig. 1: yellow, catalytic domain; purple, CDP-AP signature motif; grey, dimerization domain; brown, N-terminal stretch; cyan, phospholipid (PC) at the dimer interface: red, CDP-choline; salmon, PC in the substrate-binding site. The CDP-choline-binding pocket and the PC binding pocket are enlarged in zoom-in views showing stick models of these substrates. Part of CDP-choline is held in place by the CDP-binding motif (purple) located between TM helices 2 and 3. Two magnesium ions are present in the same place as DAG-bound state and depicted as green balls. **c.** PC binding mode analyzed by LigPlot2. His118, that exhibits alternate conformations, is colored in purple. The residue with polar interaction with PC (Ser158) is shown as ball and sticks and labeled in green. **d.** CDP-choline-binding mode viewed from the lumenal side. **e.** CDP-choline-binding residues analyzed by LigPlot2. The residues with polar interaction with CDP-choline are shown as ball and sticks and labeled in green. **f.** Displacement of the conserved tryptophan residue in the juxtamembrane helix (JMH) comparing the cryo-EM structure of yCPT1 (pale red) and the AF2 model of yEPT1 (turqouise). A conserved tryptophan (Trp35) is facing either alanine or glycine, respectively. **g, h.** Choline- or Ethanolamine-phosphotransferase activity of yCPT1 WT and yCPT1 A97G mutant was measured using CDP-choline or CDP-ethanolamine and deuterium-labeled DAG as the substrates. **i.** Time course for the PE synthesis by yCPT1 WT and yCPT1 A97G. Data are presented as the mean ± SD. *** indicates P < 0.001 as compared with yCPT1 WT. Statistical significance was determined using a Student’s t-test.

Our structure shows that yCPT1 binds the CDP part of CDP-choline with contributions from all the residues in the conserved CDP-AP signature motif (^96^**<u>D</u>**AC**<u>DG</u>**MH**<u>AR</u>**RTGQQGPL**<u>G</u>**ELF**<u>D</u>**HCI**<u>D</u>**^121^) contributing (Fig. 3b,d, and e), very similar to what was seen in other CDP-APs ^17–22,24,25^. In this binding position, the cytosine head of CDP is in the proper orientation to be fixed by a hydrogen bond network between the 4-amino group of cytosine and side chains from Asn49 and Thr52, with the pyrimidine ring sandwiched between Gly100, Ala103, and Gly113 (Fig. 3e). The ribose group is held in place by forming hydrogen bonds between the two hydroxyl groups and Tyr16 and Glu114 (Fig. 3e). The α-phosphate makes an ionic interaction with Arg104 (Fig. 3e). This ionic interaction is conserved among all reported structures of CDP-APs and is likely critical for the catalytic reaction. Mutation of the equivalent arginine in hEPT1 (R112) was reported to be associated with a complex form of hereditary spastic paraplegia (HSP) that includes intellectual impairment, demyelination, and motor function deficiency^32^. The diphosphate group coordinates one of the Mg^2+^ ions together with Asp96, Asp99, and Asp117. The choline moiety points towards the JMH facing residues Ser27 and Trp35 of this helix (Fig. 3d,e). Previous reports showed that yCPT1 is specific for CDP-choline, while yEPT1 is more promiscuous for both CDP-choline and CDP-ethanolamine^11,14–16^. Of the residues interacting with the choline moiety, Trp35 faces toward Ala97 of TM helix 2. In yEPT1, based on sequence alignment, the equivalent residue is Gly97 (Supplementary Fig. 10a). When superposing the AF2-predicted yEPT1 model on our yCPT1 cryo-EM structure, this alanine to glycine substitution in yEPT1 relocates Trp35 (equivalent to Trp35 in yCPT1) inward by up to 2.2Å (Fig. 3f). Consequently, the entire JMH also moves inward in the predicted yEPT1 structure. We initially considered the possibility of whether the position of Trp35 in yCPT1 is an induced fit by the presence of the substrate. However, the position of this residue is the same in the structure of yCPT1 without CDP-choline, indicating that its position is defined by the presence of the Ala97 side chain and not influenced by the presence of CDP-choline (Supplementary Fig. 10b). Interestingly, a previous study found that CDP-propanolamine, which differs from CDP-ethanolamine by one methylene group in the head group, is preferred by yCPT1 rather than yEPT1^16^ (Supplementary Fig. 10c). When combined with our structural model this preference suggests a correlation between the position of Trp35 in the JMH and a size preference of the head group of the substrate. To test this model, we made an alanine to glycine mutation at this site in yCPT1 (yCPT1 A97G) and investigated the substrate preference of the wild type and mutant yCPT1 by evaluating their abilities to synthesize PC or PE. Our analysis coincides with previously reports^11,14–16^; yCPT1 is selective for CDP-choline; however, yCPT1 A97G is permissive toward CDP-ethanolamine (Fig. 3g-i). Based on the previous reports and our results, we propose that the position of Trp35, or the equivalent residue in homologs, that faces towards the substrate binding pocket from the JMH defines the head-group selectivity and that this position is determined by a structurally opposing residue from TM2 such as Ala97 in yCPT1. Sequence alignment of human and yeast CDP-APs shows that hCHPT1 and hCEPT1 have an alanine at an equivalent position to Ala97 in yCPT1 and that hEPT1 has a glycine residue at this position (Supplementary Fig. 10a). Functionally, hCEPT1 is a promiscuous enzyme capable of using both CDP-choline and CDP-ethanolamine with preference towards CDP-choline, while hCHPT1 and hEPT1 are exclusive for CDP-choline or CDP-ethanolamine, respectively (Supplementary Fig. 1)^10,16,33^. Since our mutational analysis showed that in yCPT1 changing Ala97 to Gly altered substrate specificity from choline to ethanolamine, we wondered whether the identity of the amino acid at this position in other CDP-APs could predict substrate specificity. This model does seem to accurately predict hCHPT1’s and hEPT1’s substrate specificity since each enzyme respectively has either an Ala or Gly at the equivalent position of yCPT1 Ala97. However, this prediction does not work for hEPT1, which with an Ala at this position would be predicted to be specific for CDP-choline, but instead is a promiscuous enzyme capable of using both CDP-choline and CDP-ethanolamine. This indicates that there are likely additional structural features that contribute to head-group selectivity.

## Selective inhibition of yCPT1 by chelerythrine and mode of inhibition

A previous report identified that R59949 and chelerythrine act as inhibitors of hCEPT1 at mid- to high micromolar concentrations^26^, although the mechanisms of this inhibition were not explored. To carefully investigate how these compounds alter hCEPT1 and yCPT1 activity, we first tested their effect on PC synthesis using recombinant hCEPT1 and yCPT1. In these assays we observed that chelerythrine inhibited yCPT1 activity at 100 µM, but R59949 had no effect (Supplementary Fig 11a). Unexpectedly, we found that neither R59949 nor chelerythrine inhibited hCEPT1 (Supplementary Fig. 11a). Titration of chelerythrine with yCPT1 showed an IC50 of 0.724 nM to for this inhibitor (Fig. 4a and Supplementary Fig. 11b). To gain structural insights into the inhibition mechanism of chelerythrine towards yCPT1, we determined a 3.1 Å resolution cryo-EM map of yCPT1 in the presence of 10 µM chelerythrine (Fig. 4b,c, Supplementary Fig. 12, and Supplementary Table1). The structure of the yCPT1 bound to chelerythrine had a similar overall architecture to the yCPT1 DAG-bound structure (r.m.s.d. = 0.558). A density for chelerythrine in the yCPT1 map was found in the hydrophobic pocket of each monomer and its position overlaps with the channels that accommodate the DAG acyl-chains (Figs. 2-b-d and 4d). The chelerythrine benzodioxolo group binds deeply in channel 1 which normally accommodates a DAG acyl chain. The main phenanthridine body of chelerythrine is found positioned in the second acyl chain channel (channel 2) and is surrounded by several hydrophobic residues (Fig. 4d,e). Notably, the iminium moiety of chelerythrine is in the correct position of the map to form an ionic interaction with the potentially deprotonated C225 (Fig. 4d,e). Thus, it is evident from the cryo-EM structure that chelerythrine competes for the DAG acyl-chain binding sites in yCPT (Supplementary Fig. 11c). Notably, the side-chain residues surrounding chelerythrine in yCPT1 are not well conserved in hCEPT1, accounting for the insensitivity of hCEPT1 to this compound in our assay (Supplementary Fig. 11d).

**Fig. 4.**
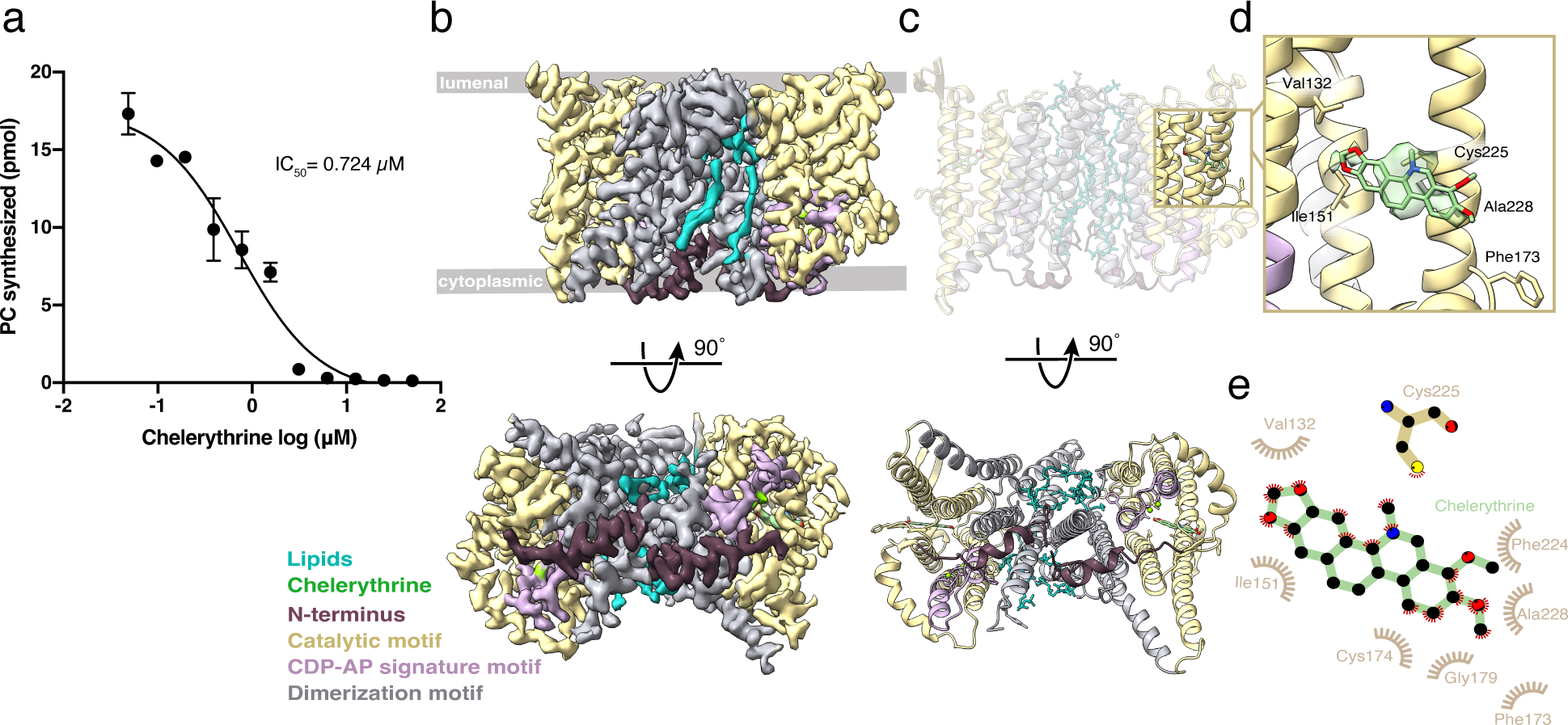
Chelerythrine is a yCPT1 inhibitor. **a.** Titration of chelerythrine for the inhibits PC synthesis activity of purified yCPT1. The data points are the mean ± SD obtained from three independent experiments. **b, c.** 3.2 Å resolution cryo-EM density map and model of yCPT1 dimer bound to chelerythrine. Density and model are depicted and colored in the same way as shown in Fig. 1: yellow, catalytic domain; purple, CDP-AP signature motif; grey, dimerization domain; brown, N-terminal stretch; cyan, phosphatidylcholine (PC). Chelerythrine is colored pale green. **d.** The chelerythrine-binding pocket in yCPT1 is enlarged with chelerythrine and surrounding residues shown as a stick models. **e.** LigPlot2 is used for the analysis of the chelerythrine-binding pocket and the illustration.

## Discussion

From our structural and function analysis of yCPT1 in multiple states, we can now propose a model for the catalytic cycle of CDP-choline and DAG conjugation by yCPT1. Within the CDP-AP signature motif, a histidine residue exhibits variable rotameric conformations across these states. In the CDP-choline bound structure, the density for His118 shows two rotameric states, one is pointing away from the catalytic site and the other is positioned towards the pyrophosphate group of CDP-choline (Supplementary Fig. 13a,b, Fig. 5a). These two rotameric states of His118 are also present in the DAG bound yCPT1 map although the density for the His118 positioned towards the pyrophosphate group of CDP-choline is weaker than the density of the other rotameric conformer. This could be due to the lower resolution of this map than the CDP-choline bound state and/or conformational heterogeneity caused by the absence of CDP-choline (Supplementary Fig. 13c). The chelerythrine-bound yCPT1 structure, since it is not bound to either CDP-choline or DAG, represents the apo-state with respect to the substrate-binding pocket. In this apo-state, His118 is positioned away from the catalytic site showing no density for the conformation of His118 positioned towards the pyrophosphate group of CDP-choline (Supplementary Fig. 13d). Together with Asp121, a part of the conserved CDP-AP signature motif, the His118 that points towards the catalytic site in the CDP-choline bound structure coordinates a water molecule between the two residues. This His118 is also hydrogen bonded with Glu114 from the opposite side of the water molecule (Fig. 5a). In xlCHPT1, the mutation of the equivalent residues abolishes enzymatic activity^25^. In line with this result, mutagenesis of His118 and Glu114 in yCPT1 abrogated the catalytic activity of yCPT1, demonstrating the functional significance of these residues (Fig. 5b). Although the coordinated water molecule would not be in the optimal position for the nucleophilic attack on the β-phosphate of the CDP-choline, since they would be > 5Å apart in the structure, it is possible that this water represents the position that the 3-hydroxyl group of the DAG would occupy prior to the attack on the β-phosphate of the CDP-choline. On the basis of the conformational variation of the amino acid side chains in this region seen between the substrate bound and the apo states of yCPT1 (Supplementary Fig. 13b-d), we propose that the binding of substrate induces the rotameric change of His118, causing it to be aligned with Tyr92, Asp121, 3-hydroxyl of DAG, Glu114, and Tyr16. And this alignment primes the deprotonation of the 3-hydroxyl of DAG by His118 and the stabilization of the protonated His118 by Glu114 to initiate the nucleophilic attack on β-phosphate (Fig. 5c). This proposed reaction mechanism is reminiscent of those seen in the catalytic sites of other lipid transferases^34–36^ and phospholipases^37^, as well as the serine proteases wherein “Asp/Glu-His” mediates the proteolytic reaction through a series of proton transfer reactions^38,39^.

**Fig. 5.**
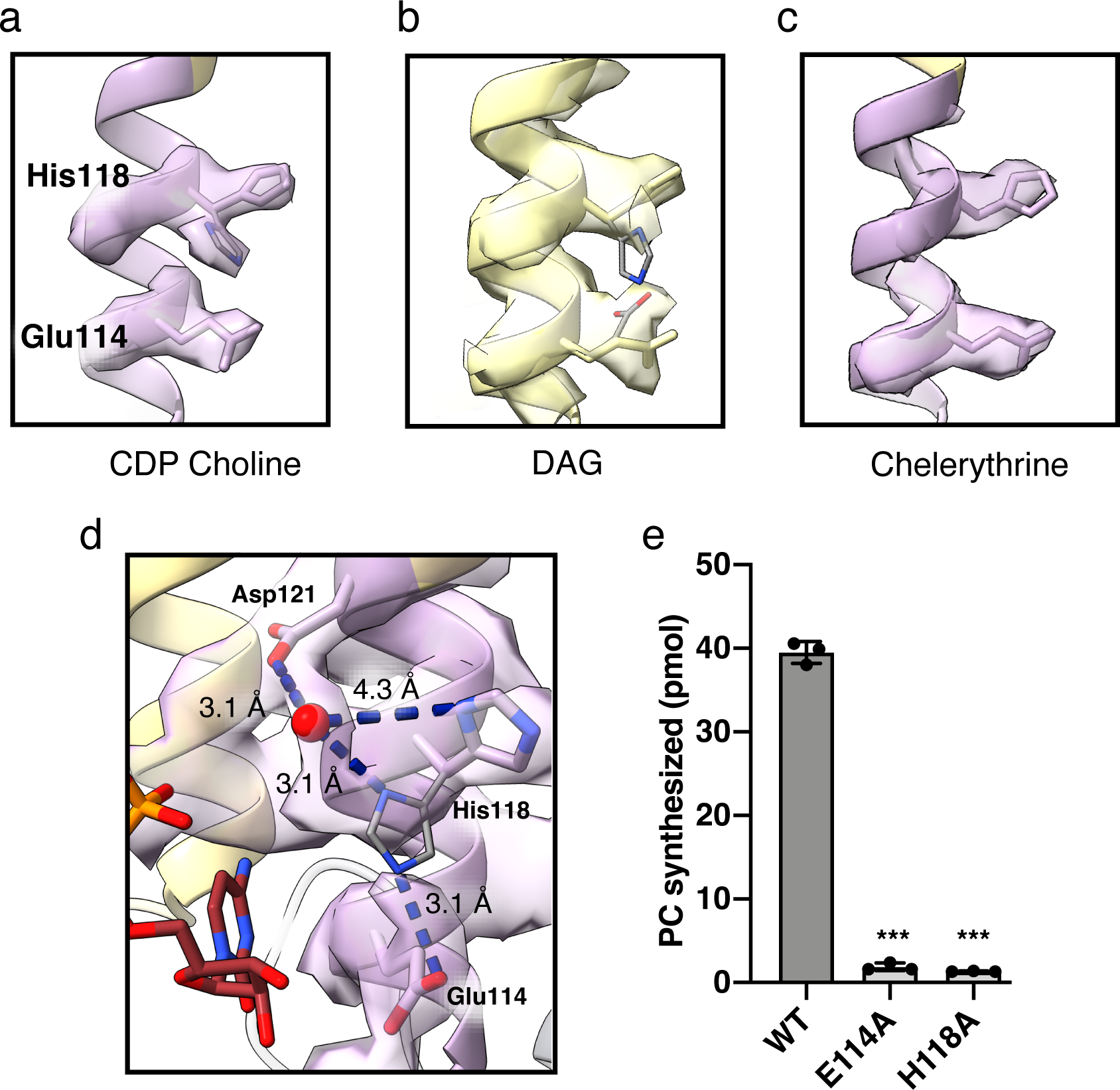
Rotameric Histidine switch and proposed catalytic mechanism of yCPT1. **a.** Structural organization and the cryo-EM density map of the rotameric His118 conformations, water molecule, Glu114, Asp121, and CDP-choline. CDP-choline, Glu114, Asp121, and His118 are depicted as sticks and water molecule is shown as a red ball. **b.** PC synthesis activity of yCPT1 WT and mutants. Data are presented as the mean ± SD. **c**. (1) In the apo-state, His118 mostly adopts an “out” conformation pointing away from the catalytic center. (2) Upon the binding of DAG, the equilibrium of His118 moves toward “in” conformation closer to the catalytic center. (3) The binding of CDP-choline further stabilizes the equilibrium of His118 to the “in” position (4) His118 at the “in” position deprotonates 3-hydroxyl of DAG to trigger nucleophilic substitution reaction toward the phosphate group of CDP-choline. (5) End-products PC and CMP are released from their binding sites and yCPT1 returns to the apo-state.

In summary, through structural, mutational, and enzymological studies, we revealed the overall architecture and the molecular basis of the substrate specificity of yCPT1, the terminal enzyme in the Kennedy pathway that produces PC. The cryo-EM structures of yCPT1 presented here reveal a unique dimeric organization compared to the structures of other CDP-AP family members^19–25^. From the structure of yCPT1 in complex with DAG we determined that a single amino acid residue (Phe146) at the exit of channel 1 is responsible for the preference in acyl-chain length. From the structure of yCPT1 in complex with CDP-choline we determined the molecular mechanism of substrate selectivity between choline and ethanolamine, which is defined by two amino acid residues (Trp35 and Ala97) in the JMH and TM2. The structure of yCPT1 in complex with chelerythrine revealed the selective inhibition mechanism of yCPT1 by this compound. These structures together allow us to propose a model for the catalytic cycle of yCPT1, where a rotameric histidine residue (His118) acts as a molecular switch. Given the ubiquitous presence and fundamental roles of phospholipids as building blocks for biological membranes, further studies will be required to apply these mechanistic insights to CDP-APs of other species.

## Acknowledgements

This work was supported by the University of Michigan, JSPS KAKENHI Grant Number 21K06824 (Y.H.), DGE 1841052 (J.R.R.) and NIH grants S10OD030275 (M.D.O.), and GM149539 to (G.G.T.). We would like to thank Megan Saab and Dr. Louise Chang for the initial help with electron microscopy. We thank Sarah Connolly, Dr. Katarina Meze, and Dr. Alexandrea Rizo for their support and discussions. We also thank Brian K. Kobilka for the laboratory support. The UM cryo-EM facility is supported by the UM Biosciences Initiative, the Beckman Foundation, and the Life Sciences Institute.

## Author Contributions

S.M. conceived the project. S.M., B.W., and A.H. cloned, expressed, and purified proteins. S.M. prepared the cryo-EM samples and collected the cryo-EM data with support from J.R.R., V.L., and A.M.R. J.R.R. led the cryoEM data processing with help from M.D.O. and S.M. S.M. modeled, refined, and analyzed the atomic structures. Y.H. carried out mass-spectroscopy analysis. Y.H. and F.E.K. carried out WT and mutant enzyme assays under the supervision of H.S. and G.G.T. J.R.R., Y.H., and S.M. prepared the manuscript under the guidance of M.D.O. and with input from all authors. M.D.O. and S.M. supervised the project.

## Materials and Methods

### Protein expression and purification

Yeast CPT1 (UniProt ID: P17898) cDNA was cloned into a pFastBac1 vector with a C-terminal Protein C tag and poly-His tag. Baculovirus was produced according to the manual provided by the manufacturer. Spodoptera frugiperda (Sf9) cells were infected with the amplified virus at density of 3.0 to 4.0 × 10^6^ cells per ml. After 48 h incubation at 27 °C at 130rpm, cells were harvested by centrifugation and dounce-homogenized in a lysis buffer containing 20mM Tris pH7.5, 150mM NaCl, 160ug/ml Benzamidine, 2.5uM Leupeptin. Upon homogenization, the NaCl concentration was increased to 650mM and further homogenized several times. The lysis (typically less than 400mL total for 4L culture) was centrifuged at 131,000g for 60 min to collect the membrane fraction. The membrane was solubilized by dounce-homogenization with a solubilization buffer of 20mM HEPES pH7.5, 150mM NaCl, 160ug/ml benzamidin, 25ug/ml Leupeptin, 1% n-dodecyl-b-D-maltoside (DDM, Anatrace), 0.1% Lauryl maltose neopentylglycol (MNG, Anatrace), 0.21% cholesteryl hemisuccinate (CHS, Steraloids) and incubated at 4 °C for 2 h. After centrifugation at 131,000 *g* for 60 min, the supernatant was incubated with cobalt resin (Thermo Fisher Scientific) at 4 °C for 1.5 h. The resin was rinsed with a high-salt wash buffer containing 20mM HEPES pH7.5, 300mM NaCl, 0.05% DDM, 0.01% MNG, 0.011% CHS, 10mM imidazole followed by a low-salt wash buffer of 20mM HEPES pH7.5, 150mM NaCl, 0.01% MNG, 0.001% CHS, 10mM imidazole. Protein was eluted with an elution buffer containing 20mM HEPES pH7.5, 150mM NaCl, 0.005% MNG, 0.0005% CHS, 250mM imidazole. Elution was concentrated and further purified by Superdex 200 Increase 10/300 GL column (Cytiva) in a buffer containing 20mM HEPES pH7.5, 150mM NaCl, 0.005% MNG, 0.0005% CHS, 10mM MgCl_2_. Peak fractions were pooled, concentrated to 8.2mg/ml with 10% glycerol, and flash frozen in small aliquots for the downstream experiments. All yCPT mutants were generated using the QuikChange method (Stratagene) and the entire cDNA was sequenced to verify the mutation. Mutants were expressed and purified following the same protocol as wild type yCPT.

Nanobodies against purified yCPT were generated using a synthetic nanobody library^27^. For the first round of selection, approximately 5x 10^9^ yeast cells were selected with 1 μM of purified yCPT in 20mL. After incubation at 4 °C for 1.5 h in a selection buffer (20 mM HEPES pH 7.5, 150 mM NaCl, 0.05% MNG, 0.005% CHS, 0.1% w/v bovine serum albumin, 5 mM CaCl_2_ and 5 mM maltose), yeast cells were washed twice with ice cold selection buffer and incubated with FITC-labelled anti-Protein C tag antibody for 15 min at 4 °C. Subsequently, yeast cells were washed twice and incubated for 15 min with anti-FITC microbeads (Miltenyi Biotec) at 4 °C. Finally, yeast cells bound with yCPT were enriched by magnetic affinity cell sorting (MACS, Miltenyi Biotec). Recovered yeast cells were passaged into SDCAA medium and induced again in SGCAA medium according to the manual previously described. Prior to the second round of selection, a negative selection was performed to deplete nanobodies that bind to the Alexa647 fluorophore or the anti-Protein C tag antibody. Approximately 2x 10^8^ yeast cells were first incubated with Alexa647-labelled anti-Protein C antibody for 15 min 4 °C, washed twice, incubated with anti-Cy5 microbeads (Miltenyi Biotec), and loaded through a MACS LD column. Unbound yeast population was subsequently used for the positive selection in the same procedure as the first round with 1 μM yCPT in 3mL. The final two rounds of selection used fluorescence-activated cell sorting (FACS). Yeast cells were first negatively selected against Alexa647 fluorophore-labeled anti-Protein C tag antibody in the same way as the second round. Unbound yeast cells were incubated with 1 μM yCPT for 15 min at 4 °C, then washed twice with selection buffer, and subsequently incubated with Alexa647-labelled anti-Protein C tag antibody and Alexa488-labelled anti-HA antibody (Cell Signaling Technology) for 15 min at 4 °C. A small fraction of yeast population (0.1-0.2% of total clones) with high signal in the Alexa647 channel were isolated and expanded in SDCAA medium. After induction in SGCAA medium, another round of FACS was performed in the same procedure as the third round. Post-4^th^ round yeast miniprep library was transformed into DH5alpha *Escherichia coli* and 7 unique clones were identified by the Sanger sequencing.

The sequence for each nanobody was amplified by PCR from the yeast surface display vector and cloned into pET26b (Novagen) with a C-terminal hexahistidine tag by Gibson assembly. The construct was transformed into BL21(DE3) Rosetta *E. coli,* grown in terrific broth medium supplemented with 2 mM MgCl_2_ and 0.1% w/v glucose and induced at an optical density (OD) at 600 nm of 0.6-1.0 with 0.5 mM IPTG. After induction, cells were further grown for 18-20 h at 18 °C and harvested by centrifugation. The bacterial pellet was stored at -80°C until purification. The cells were resuspended in over 25 ml of TES buffer (200 mM Tris pH 8.0, 500 mM sucrose, 0.5 mM EDTA, 0.1mM PMSF) per 15 gram of the cell. After stirring for 45 min at 4 °C, two volumes of cold MilliQ water were added to induce hypotonic lysis of the periplasm and the bacteria suspension was further stirred at 4 °C for 45 min. The resulting lysate was centrifuged at 18,000g for 15 min at 4 °C. The supernatant was supplemented with 1mM MgCl_2_ to quench the EDTA and incubated with Ni-NTA resin for 1h at 4 °C. The resin was washed extensively with a high-salt wash buffer containing 20 mM HEPES pH 7.5, 500 mM NaCl and 20 mM imidazole, and eluted with an elution buffer containing 20 mM HEPES pH 7.5, 500 mM NaCl and 300 mM imidazole. Eluted nanobody fractions were pooled and further purified by size-exclusion chromatography using Superdex 200 Increase 10/300 GL column (Cytiva) in a buffer containing 20mM HEPES pH7.5, 150mM NaCl. Peak fractions were concentrated to over 20 mg/ml, supplemented with 10% glycerol, and flash frozen in small aliquots in liquid nitrogen and stored at −80 °C until further use.

### Sample preparation and data acquisition for cryo-EM

To prepare the cryo-EM sample, yCPT protein was incubated with over 2-fold excess nanobody24 (Nb24) for 60 min at 4 °C. Excess Nb24 was removed by size-exclusion chromatography (SEC) using Superdex 200 Increase 10/300 GL column (Cytiva) in a buffer containing 20mM HEPES pH7.5, 150mM NaCl, 0.005% MNG, 0.0005% CHS, 10mM MgCl_2_. Peak fractions containing both yCPT and Nb24 were concentrated using a 100-kDa molecular weight cutoff concentrator (Millipore) to 2 to 4 mg/ml. For yCPT bound with chelerythrine, yCPT-Nb25 complex was incubated with 10µM chelerythrine for 1h on ice prior to SEC and the peak fractions were again incubated with 10µM chelerythrine and concentrated to over 1mg/ml. For yCPT bound with CDP-choline, yCPT-Nb25 complex was incubated with 1mM CDP-choline for 1h at room temperature prior to SEC and the peak fractions were again incubated with 1mM CDP-choline and concentrated to over 3mg/ml. Grids (Quantifoil Cu R1.2/1.3, 400 mesh) were glow-discharged for 30 s and 3.5 µL of the protein at 1.0 to 3.0 mg/ml were applied to the grids and blotted for 3.5 s before being flash-frozen in liquid ethane cooled by liquid nitrogen with Vitrobot Mark IV (Thermo Fisher Scientific) at 4 °C and 100% humidity. The grids were stored in liquid nitrogen until cryoEM observation. The grids were transferred to a 300 kV Titan Krios equipped with Gatan K3 Summit detector and a GIF Quantum energy filter (slit width 20 eV). Micrographs were recorded at a magnification of 81,000 X corresponding to 1.08 pixel per A° at the specimen level with a defocus range from −2.4 to −1.5 μm. Each stack of 60 frames was exposed for 4.4 s, with an exposing time of 0.073 s per frame. The total dose was about 60 e^-^/Å^2^ for each stack. All image stacks were collected by SerialEM.

### CryoEM Data processing

For yCPT in complex with 1,2-diacylglycerol, 3,625 movies were patch motion corrected and patch CTF corrected in cryoSPARC v 4.2.1^40^. A representative micrograph is shown in Figure S1A. 2,913,379 particles were picked using crYOLO^41^ and extracted using a 256 pixel box size (1.08 Å/pixel). Iterative 2D classification and the removal of duplicate particles resulted in 1,651,587 particles, of which representative 2D classes are shown in Figure S1B. These particles were further processed in a four-class *ab initio* 3D reconstruction (Figure S1C). Of the four resulting volumes, three had density embedded in the detergent micelle. The 1,363,364 particles from these three structures were used in heterogenous refinement designated into two classes. The major class containing 965,534 particles exhibited clear transmembrane helices. These particles were used in an additional round of heterogeneous refinement designated into three classes. The best resolved 3D class contained 677,515 particles (Figure S1D). Subsequent non-uniform refinement with no applied symmetry (C1) leads to a structure at 3.2 Å resolution (Figure S1E).^42^ Local refinement was used to resolve ligands, such as diacylglycerol (DAG), and the lipids lining the dimer interface. Local resolution was estimated in cryoSPARC using the “Local Resolution Estimation” job (Figure S1G).

For yCPT in complex with CDP-choline, a total of 9,184 movies were patch motion corrected and patch CTF corrected in cryoSPARC v 4.2.1^40^. A representative micrograph is shown in panel S2A. 2,913,279 initial particles were picked using crYOLO^41^, extracted with a box size of 256 pixel (1.08 Å/pixel) and then Fourier cropped to 128 pixel (2.16 Å/pixel). After iterative 2D classification, 2,861,130 particles were selected. Representative 2D classes are shown in Figure S2B. An *ab initio* 3D reconstruction was done designated into 2 classes and the class containing a better defined structural features was used as the 3D reference for non-uniform refinement^42^, using the unbinned particles (box size of 256 pixels(1.08 Å/pixel)). Because of strongly preferred orientation of the particles in the vitrified ice layer (Figure S2E), additional data was collected by adding a detergent CHAPS (3-((3-cholamidopropyl) dimethylammonio)-1-propanesulfonate) to the sample to a final concentration of 3.25 mM prior to the sample vitrification. 2,522 movies were collected and processed in the same way described for the non-CHAPS sample. A representative micrograph is shown in panel S2B. crYOLO^41^ was used to pick 572,751 initial particles and extracted with a box size of 256 pixels (1.08 Å/pixel) and then Fourier cropped to 128 pixel (2.16 Å/pixel). After iterative 2D classification 75,015 good particles were selected and used in *ab initio* 3D reconstruction designated into two classes. The 3D class with a more structural features was used as the initial volume for non-uniform refinement^42^, using the unbinned box size of 256 pixels (1.08 Å/pixel). The refinement reached a resolution of 3.5 Å, (Figure S2F). The particles used in the non-uniform refinement step for plus- and non-CHAPS, were combined (1,228,251 particles), for another round of non-uniform refinement^42^ (Figure S2G) with no applied symmetry (C1) reaching a resolution of 2.96 Å (Figure S2H) with a better particle distribution (Figure S2I). Local resolution was estimated in cryoSPARC^40^ using the “Local Resolution Estimation” job (Figure S2J).

For yCPT in complex with chelerythrine, 2,990 movies were patch motion corrected and patch CTF corrected in cryoSPARC v 4.2.1^40^. Particles were picked from motion-corrected micrographs (Figure S3A) using the cryoSPARC template picker. Templates were created from 2D projections of the final volume of the apo dataset that were lowpass filtered to 20 Å. 4,613,867 particles were picked and extracted using a 256 pixel box size (1.08 Å/pixel) and then Fourier cropped to 128 pixels (2.16 Å/pixel). Iterative 2D classification and the removal of duplicate particles resulted in 1,818,356 particles, of which representative 2D classes are shown in Figure S3B. These particles were used in *ab initio* 3D reconstruction with three designated classes. Of the three *ab initio* 3D volumes, one had the clear transmembrane helical density. The 522,909 particles assigned to this volume were unbinned to 256 pixels (1.08 Å/pixel) and used in heterogenous refinement that designated into two classes. A major class with more defined structural features contained 371,370 particles. This volume and corresponding particles were used in non-uniform refinement with no applied symmetry (C1) reaching 3.2 Å resolution (Figure S3D,E). Local resolution was estimated in cryoSPARC using the “Local Resolution Estimation” job (Figure S3G).

Map and model figures were created using ChimeraX^43^.

### Structural model building and refinement

An initial structure model for yCPT was obtained by AlphaFold2^31^. The structure was docked into the density map The restraints for substrates were generated by eLBOW in the Phenix software suite^44^. Substrate incorporation into the protein chain was performed manually and with real-space refinements in Coot. Followed by iterative manual adjustment in COOT and phenix.real_space_refine in PHENIX, the final structure of yCPT was validated using Molprobity^45^. Owing to the higher resolution and better quality of the CDP-choline bound yCPT reconstruction relative to the other reconstructions, coordinates after real-space refinement against the CDP-choline bound yCPT were used as an initial model of yCPT in other states, and minor adjustments were made before running a final real_space_refinement. The molecular graphic figures were prepared with UCSF ChimeraX^46^ and Pymol.

### Sequence conservation analysis by Consurf

Conservation within the CDP-AP family was analyzed using the ConSurf server^47^. CDP-choline bound yCPT was used as an input model and non-redundant homolog sequences were retrieved from UniProt database using HMMER. Multiple sequence alignment was built using MAFFT. The result from the server is visualized with PyMol.

### Enzyme assay

Enzymatic activity of choline phosphotransferase was measured as previously described^48^. DAG 16:0–16:0 and DAG 18:1–18:1 were obtained from Sigma-Aldrich and Avanti Polar Lipids, respectively. The reaction mixture contained 20 mM Tris-HCl buffer, pH 8.0, 30 mM MgCl_2_, 0.003% Tween 20 (w/v), 10 μM radiolabeled CDP-choline (1,2 -^14^C-CDP-choline, American Radiolabeled Chemicals), 200 μM DAG, and enzyme. After incubation at 37°C for 5 min, radiolabeled PC was separated by to a TLC plate (Merck), which was developed with chloroform-methanol-water (65:38:8, v/v/v). Radiolabeled PC were analyzed using an FLA-7000 imaging analyzer and quantified using ImageQuant TL version 8.1 (GE Healthcare). For ethanolamine phosphotransferase assay, enzyme was incubated in 20 mM Tris-HCl buffer, pH 8.0, 30 mM MgCl_2_, 0.003% Tween 20 (w/v), 20 μM CDP-ethanolamine (Sigma-Aldrich), 200 μM deuterium-labeled DAG (DAG 15:0–18:1-d7, Avanti Polar Lipids). After incubation at 37°C for 5 min, the deuterium-labeled PE was quantified by LC-MS/MS as described below.

### Extraction and quantification of lipid by LC-MS/MS

The lipid was extracted from the purified yCPT1 (900 pmol) with methanol. The levels of PC, PE, and DAG were measured by LC-MS/MS as described previously^48^. The lipid was analyzed by reverse-phase HPLC using an L-column 3 ODS column (3 µm, 2.0 × 100 mm) (Chemicals Evaluation and Research Institute, Tokyo, Japan) coupled to a 5500 QTRAP mass spectrometer (Sciex Inc.). PC and PE were detected in multiple reaction monitoring mode by selecting the m/z of the lipid molecular species at Q1 and the precursor of m/z 184 for PC and neutral loss of m/z 141 for PE at Q3, respectively, in positive ion mode. DAG was detected as described previously^49^. The deuterium-labeled PE was detected in positive ion mode with m/z transitions at 711.5/570.5. Peak area of lipid was quantified using MultiQuant version 2.0 (Sciex).

### Statistical analysis

Quantitative data are presented as means ± S.D. Statistical significance was assessed using a Student’s t-test or a one-way ANOVA with Tukey’s post hoc test. A p-value of <0.05 was considered statistically significant.

## References

1. van Meer, G., Voelker, D. R. & Feigenson, G. W. Membrane lipids: where they are and how they behave. Nat. Rev. Mol. Cell Biol. 9, 112–124 (2008).

2. Saliba, A.-E., Vonkova, I. & Gavin, A.-C. The systematic analysis of protein–lipid interactions comes of age. Nat. Rev. Mol. Cell Biol. 16, 753–761 (2015).

3. Vance, J. E. Phospholipid Synthesis and Transport in Mammalian Cells. Traffic 16, 1–18 (2015).

4. Shimizu, T. Lipid Mediators in Health and Disease: Enzymes and Receptors as Therapeutic Targets for the Regulation of Immunity and Inflammation. Annu. Rev. Pharmacol. Toxicol. 49, 123–150 (2009).

5. Lamari, F., Mochel, F., Sedel, F. & Saudubray, J. M. Disorders of phospholipids, sphingolipids and fatty acids biosynthesis: toward a new category of inherited metabolic diseases. J. Inherit. Metab. Dis. 36, 411–425 (2013).

6. Gibellini, F. & Smith, T. K. The Kennedy Pathway-De Novo Synthesis of Phosphatidylethanolamine and Phosphatidylcholine. 62, 414–428 (2010).

7. Kennedy, E. P. & Weiss, S. B. THE FUNCTION OF CYTIDINE COENZYMES IN THE BIOSYNTHESIS OF PHOSPHOLIPIDES. J. Biol. Chem. 222, 193–214 (1956).

8. Kent, C. Eukaryotic Phospholipid Biosynthesis. Annu. Rev. Biochem. 64, 315–343 (1995).

9. McMaster, C. R. & Bell, R. M. Phosphatidylcholine biosynthesis via the CDP-choline pathway in Saccharomyces cerevisiae. Multiple mechanisms of regulation. J. Biol. Chem. 269, 14776–14783 (1994).

10. Henneberry, A. L., Wistow, G. & McMaster, C. R. Cloning, Genomic Organization, and Characterization of a Human Cholinephosphotransferase *. J. Biol. Chem. 275, 29808–29815 (2000).

11. McMaster, C. R. & Bell, R. M. Phosphatidylcholine biosynthesis in Saccharomyces cerevisiae. Regulatory insights from studies employing null and chimeric sn-1,2-diacylglycerol choline- and ethanolaminephosphotransferases. J. Biol. Chem. 269, 28010–28016 (1994).

12. Henneberry, A. L. & McMaster, C. R. Cloning and expression of a human choline/ethanolaminephosphotransferase: synthesis of phosphatidylcholine and phosphatidylethanolamine. Biochem. J. 339, 291–298 (1999).

13. Williams, J. G. & McMaster, C. R. Scanning Alanine Mutagenesis of the CDP-alcohol Phosphotransferase Motif of Saccharomyces cerevisiaeCholinephosphotransferase*. J. Biol. Chem. 273, 13482–13487 (1998).

14. Hjelmstad, R. H., Morash, S. C., McMaster, C. R. & Bell, R. M. Chimeric enzymes. Structure-function analysis of segments of sn-1,2-diacylglycerol choline- and ethanolaminephosphotransferases. J. Biol. Chem. 269, 20995–21002 (1994).

15. Hjelmstad, R. H. & Bell, R. M. sn-1,2-diacylglycerol choline- and ethanolaminephosphotransferases in Saccharomyces cerevisiae: Mixed micellar analysis of the CPT1 and EPT1 gene products*. J. Biol. Chem. 266, 4357–4365 (1991).

16. Boumann, H. A., de Kruijff, B., Heck, A. J. R. & de Kroon, A. I. P. M. The selective utilization of substrates in vivo by the phosphatidylethanolamine and phosphatidylcholine biosynthetic enzymes Ept1p and Cpt1p in yeast. FEBS Lett. 569, 173–177 (2004).

17. Grāve, K., Bennett, M. D. & Högbom, M. Structure of Mycobacterium tuberculosis phosphatidylinositol phosphate synthase reveals mechanism of substrate binding and metal catalysis. Commun. Biol. 2, 1–11 (2019).

18. Belcher Dufrisne, M., et al. Structural and Functional Characterization of Phosphatidylinositol-Phosphate Biosynthesis in Mycobacteria. J. Mol. Biol. 432, 5137–5151 (2020).

19. Clarke, O. B. et al. Structural basis for phosphatidylinositol-phosphate biosynthesis. Nat. Commun. 6, 8505 (2015).

20. Centola, M., van Pee, K., Betz, H. & Yildiz, Ö. Crystal structures of phosphatidyl serine synthase PSS reveal the catalytic mechanism of CDP-DAG alcohol O-phosphatidyl transferases. Nat. Commun. 12, 6982 (2021).

21. Yang, B., Yao, H., Li, D. & Liu, Z. The phosphatidylglycerol phosphate synthase PgsA utilizes a trifurcated amphipathic cavity for catalysis at the membrane-cytosol interface. Curr. Res. Struct. Biol. 3, 312–323 (2021).

22. Sciara, G. et al. Structural basis for catalysis in a CDP-alcohol phosphotransferase. Nat. Commun. 5, 4068 (2014).

23. Nogly, P. et al. X-ray structure of a CDP-alcohol phosphatidyltransferase membrane enzyme and insights into its catalytic mechanism. Nat. Commun. 5, 4169 (2014).

24. Wang, Z., Yang, M., Yang, Y., He, Y. & Qian, H. Structural basis for catalysis of human choline/ethanolamine phosphotransferase 1. Nat. Commun. 14, 2529 (2023).

25. Wang, L. & Zhou, M. Structure of a eukaryotic cholinephosphotransferase-1 reveals mechanisms of substrate recognition and catalysis. Nat. Commun. 14, 2753 (2023).

26. Wright, M. M. & McMaster, C. R. PC and PE synthesis: Mixed micellar analysis of the cholinephosphotransferase and ethanolaminephosphotransferase activities of human choline/ethanolamine phosphotransferase 1 (CEPT1). Lipids 37, 663–672 (2002).

27. McMahon, C. et al. Yeast surface display platform for rapid discovery of conformationally selective nanobodies. Nat. Struct. Mol. Biol. 25, 289–296 (2018).

28. McMaster, C. R., Morash, S. C. & Bell, R. M. Phospholipid and cation activation of chimaeric choline/ethanolamine phosphotransferases. Biochem. J. 313, 729–735 (1996).

29. Hjelmstad, R. H. & Bell, R. M. [32] choline- and ethanolaminephosphotransferases from Saccharomyces cerevisiae. in Methods in Enzymology (eds. Dennis, E. A. & Vance, ennis E.) vol. 209 272–279 (Academic Press, 1992).

30. Krissinel, E. & Henrick, K. Inference of Macromolecular Assemblies from Crystalline State. J. Mol. Biol. 372, 774–797 (2007).

31. Jumper, J. et al. Highly accurate protein structure prediction with AlphaFold. Nature 1–11 (2021) doi:10.1038/s41586-021-03819-2.

32. Ahmed, M. Y. et al. A mutation of EPT1 (SELENOI) underlies a new disorder of Kennedy pathway phospholipid biosynthesis. Brain 140, 547–554 (2017).

33. Horibata, Y. & Hirabayashi, Y. Identification and characterization of human ethanolaminephosphotransferase1. J. Lipid Res. 48, 503–508 (2007).

34. Rana, M. S. et al. Fatty acyl recognition and transfer by an integral membrane S-acyltransferase. Science 359, eaao6326 (2018).

35. Wang, L. et al. Structure and mechanism of human diacylglycerol O-acyltransferase 1. Nature 581, 329–332 (2020).

36. Johnson, Z. L. et al. Structural basis of the acyl-transfer mechanism of human GPAT1. Nat. Struct. Mol. Biol. 30, 22–30 (2023).

37. Selvy, P. E., Lavieri, R. R., Lindsley, C. W. & Brown, H. A. Phospholipase D: Enzymology, Functionality, and Chemical Modulation. Chem. Rev. 111, 6064–6119 (2011).

38. Bachovchin, W. W. Review: Contributions of NMR spectroscopy to the study of hydrogen bonds in serine protease active sites. Magn. Reson. Chem. 39, S199–S213 (2001).

39. Hedstrom, L. Serine Protease Mechanism and Specificity. Chem. Rev. 102, 4501–4524 (2002).

40. Punjani, A., Rubinstein, J. L., Fleet, D. J. & Brubaker, M. A. cryoSPARC: algorithms for rapid unsupervised cryo-EM structure determination. Nat. Methods 14, 290–296 (2017).

41. Wagner, T. et al. SPHIRE-crYOLO is a fast and accurate fully automated particle picker for cryo-EM. Commun. Biol. 2, 218 (2019).

42. Punjani, A., Zhang, H. & Fleet, D. J. Non-uniform refinement: adaptive regularization improves single-particle cryo-EM reconstruction. Nat. Methods 17, 1214–1221 (2020).

43. Pettersen, E. F. et al. UCSF CHIMERAX: Structure visualization for researchers, educators, and developers. Protein Sci. 30, 70–82 (2021).

44. Adams, P. D. et al. PHENIX: a comprehensive Python-based system for macromolecular structure solution. Acta Crystallogr. D Biol. Crystallogr. 66, 213–221 (2010).

45. Chen, V. B. et al. MolProbity: all-atom structure validation for macromolecular crystallography. Acta Crystallogr. D Biol. Crystallogr. 66, 12–21 (2010).

46. Goddard, T. D. et al. UCSF ChimeraX: Meeting modern challenges in visualization and analysis. Protein Sci. 27, 14–25 (2017).

47. Ashkenazy, H. et al. ConSurf 2016: an improved methodology to estimate and visualize evolutionary conservation in macromolecules. Nucleic Acids Res. 44, W344–W350 (2016).

48. Horibata, Y. & Sugimoto, H. Differential contributions of choline phosphotransferases CPT1 and CEPT1 to the biosynthesis of choline phospholipids. J. Lipid Res. 62, (2021).

49. Takeda, H. et al. Widely-targeted quantitative lipidomics method by supercritical fluid chromatography triple quadrupole mass spectrometry [S]. J. Lipid Res. 59, 1283–1293 (2018).

50. Pettersen, E. F. et al. UCSF Chimera? A visualization system for exploratory research and analysis. J. Comput. Chem. 25, 1605–1612 (2004).

51. Laskowski, R. A. & Swindells, M. B. LigPlot+: Multiple Ligand–Protein Interaction Diagrams for Drug Discovery. J. Chem. Inf. Model. 51, 2778–2786 (2011).

