## Supplemental files for "Structural basis for catalysis and selectivity of phospholipid synthesis by eukaryotic choline-phosphotransferase"

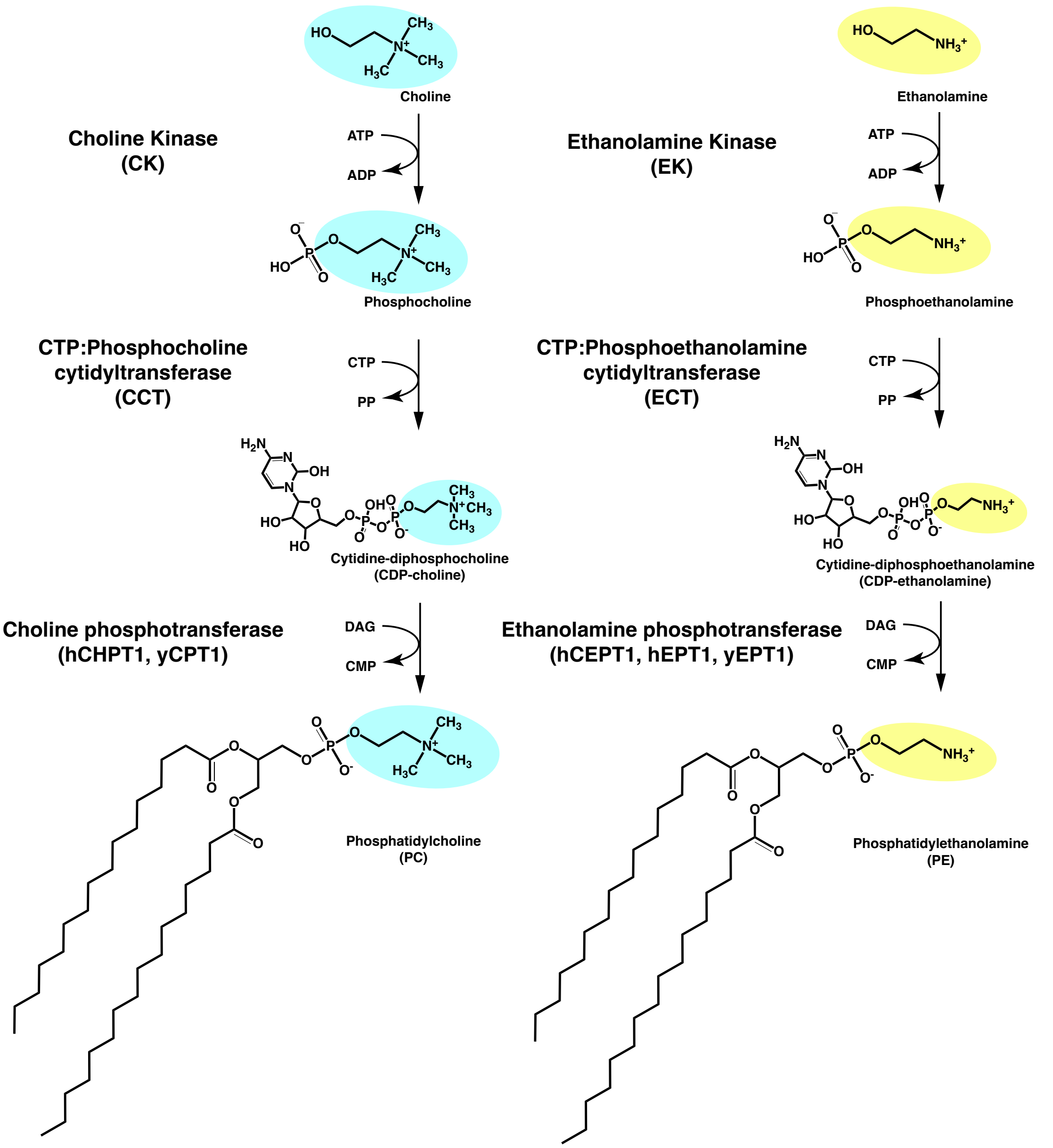

**Supplementary Figure 1. Depiction of Kennedy pathway for *de novo* phosphatidylcholine (PC) and phosphatidylethanolamine (PE) synthesis.** Choline or Ethanolamine is taken up by the cell and converted to phosphocholine or phosphoethanolamine by choline kinase or ethanolamine kinase respectively. These molecules are further converted to CDP-choline or CDP-ethanolamine by CTP:phosphocholine cytidylyltransferase or CTP:phosphoethanolamine cytidylyltransferase respectively. Choline phosphotransferase or ethanolamine phosphotransferase mediates the final step to conjugate these molecules with DAG to produce PC or PE.

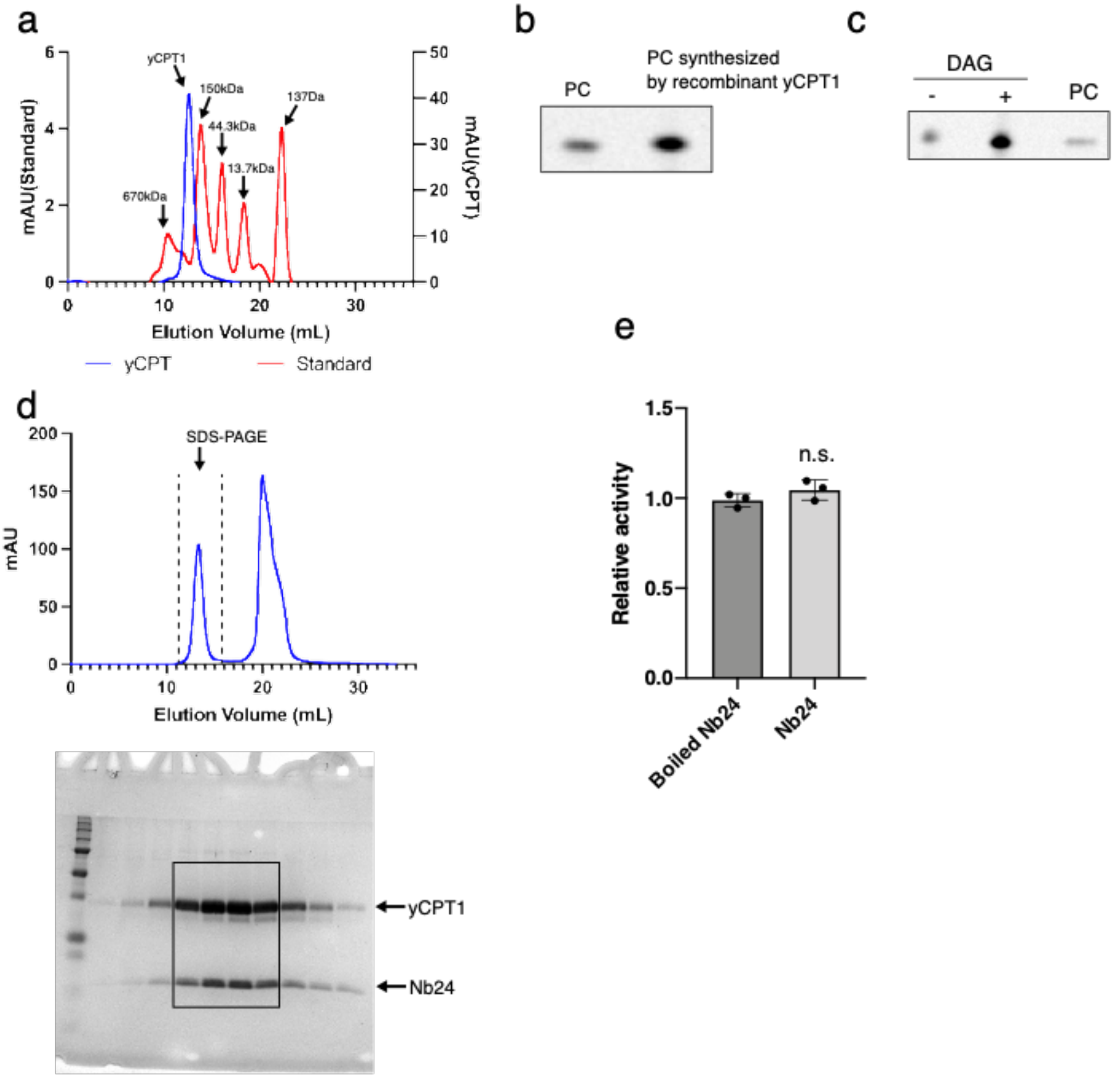

**Supplementary Figure 2. Biochemical characterization of yCPT1. a.** Analytical size-exclusion chromatography profile of yCPT1 purified from Sf9 cells overlayed with trace from molecular weight standards. **b.** The enzymatic activity of purified recombinant yCPT1 was measured using radiolabeled CDP-choline and DAG as the substrates. Radioactive phospholipids were analyzed by TLC. **c.** Purified yCPT1 contain endogenous DAG from the Sf9 cells as the enzymatic reaction proceeds without adding extra DAG. **d.** Preparative size-exclusion chromatography of the yCPT1-Nb24 complex. The main peak fractions around 12ml, shown by dashed lines, were analyzed by SDS-PAGE to confirm co-migration of Nb24. The fractions indicated by the rectangle were used to prepare the cryo-EM sample. **e.** Binding of Nb24 has no impact on the enzymatic activity of purified recombinant yCPT1. yCPT1 was preincubated with either intact Nb24 or boiled Nb24 at 100°C for 10 min prior to the reaction.

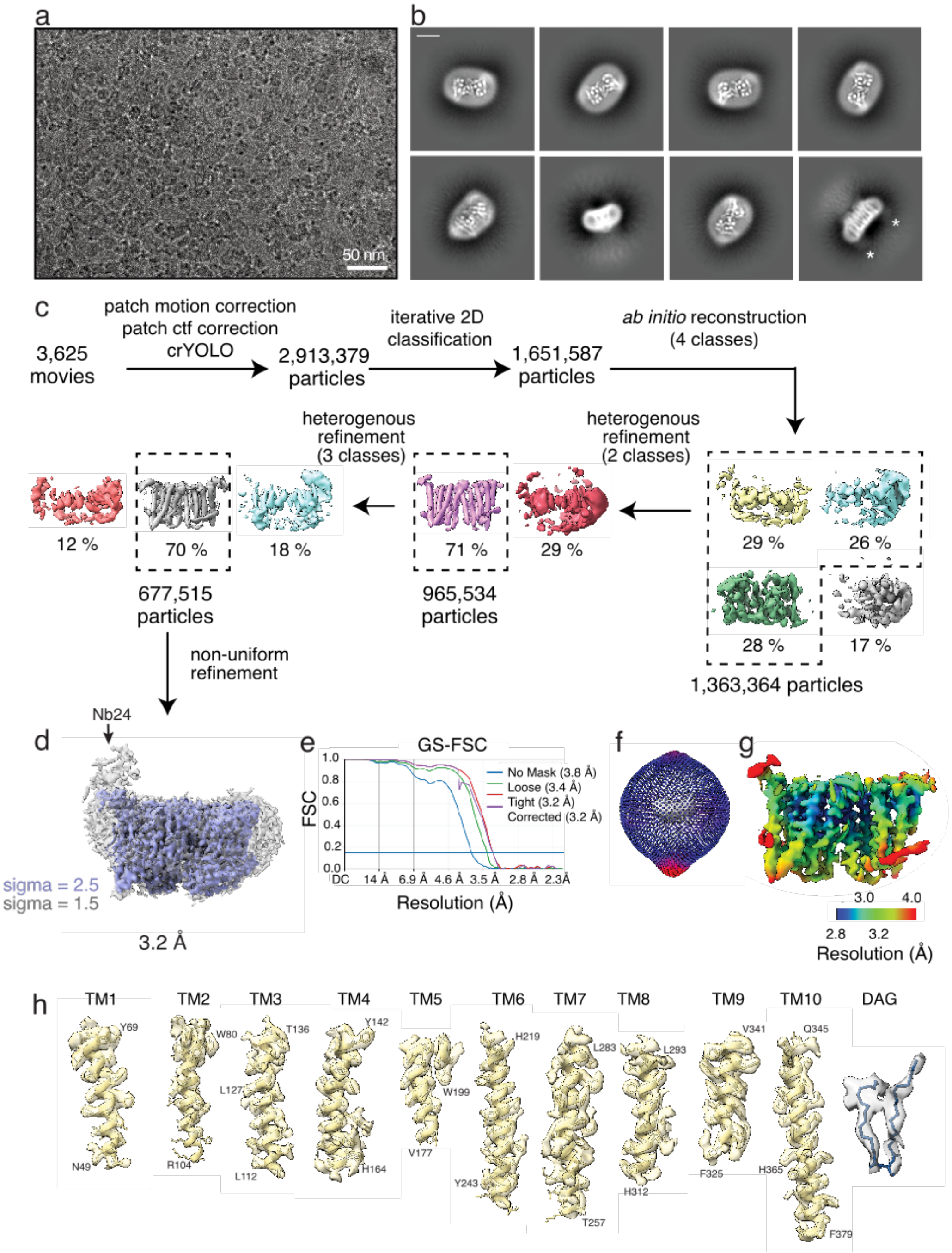

**Supplementary Figure 3. Cryo-EM processing of yCPT1-Nb24 in complex with DAG. a.** Representative sub-area of a motion corrected micrograph of vitrified yCPT1-Nb24 particles. **b.** Representative 2D classes. Position of Nb24 marked by *. Box size 276 Å. Scale bar 50 Å. **c.** Cryo-EM processing workflow. 2,913,379 particles were selected from 3,625 micrographs using crYOLO. **d.** After iterative 2D classification and heterogenous refinement 677,515 good particles were subjected to a non-uniform refinement, reaching 3.2 Å **e.** GS-FSC plot. **f**. Particle orientation distribution plot. **g.** Local resolution estimation. **h.** Density of each TM helix and DAG are shown contoured at 3 sigma.

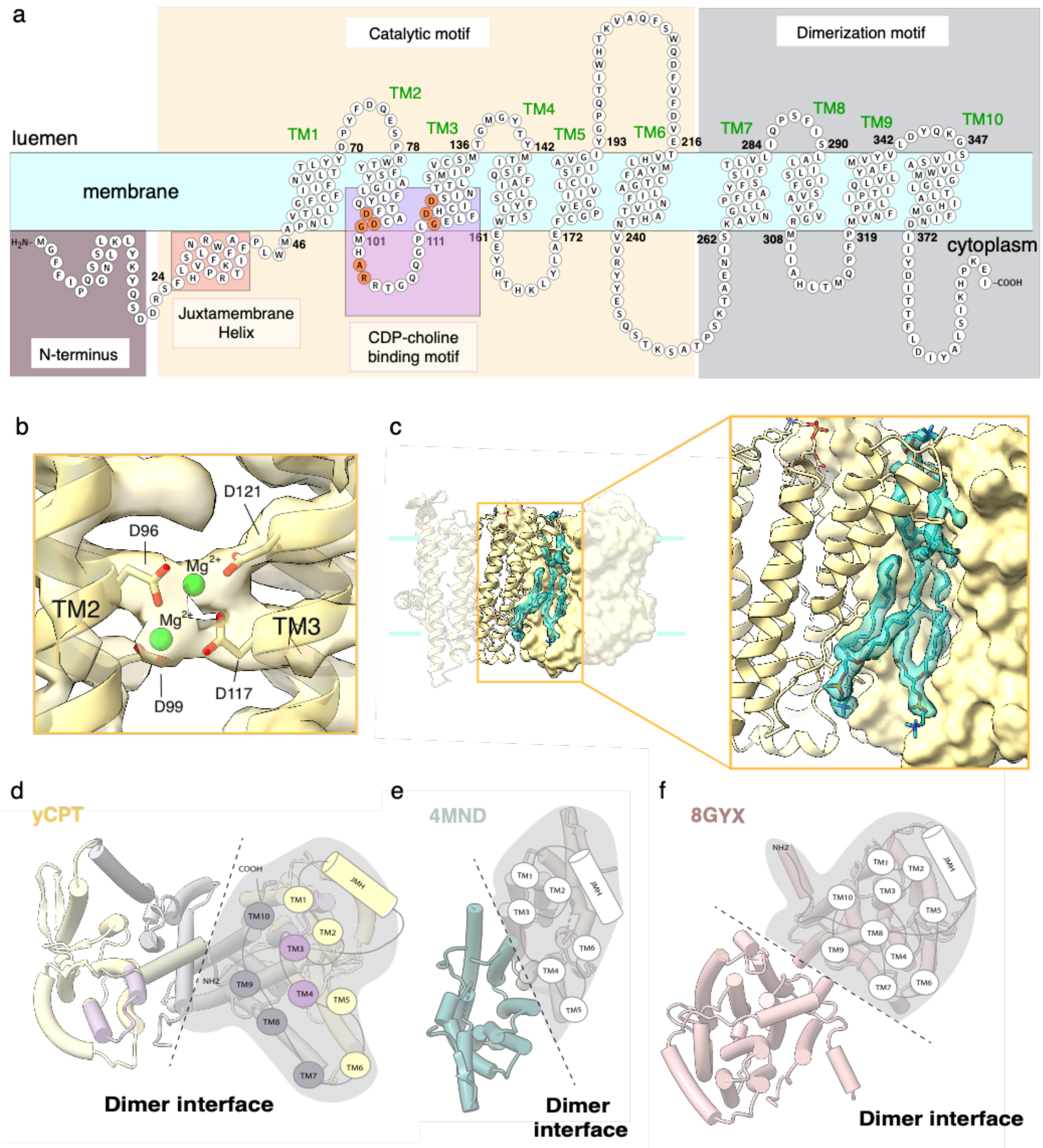

**Supplementary Figure 4. Overall architecture of yCPT1. a.** Snake plot of yCPT1 with each motif colored in the same way as in Fig.1: yellow, catalytic motif; purple, CDP binding motif; grey, dimerization motif; ruby red, N-terminal stretch. The juxtamembrane helix is represented in a red rectangle and CDP-AP signature motif residues are colored in orange. **b.** Magnesium coordination at the signature motif with the cryo-EM density map surrounding the magnesium ions. **c.** Phospholipid molecules embedded between monomers of yCPT1. Each protomer is represented as cartoon and surface, and the phospholipids are shown as sticks with the cryo-EM density. **d.** Distinct dimer interface of yCPT1 in comparison to prokaryotic or other eukaryotic CDP-APs. - *A. fulgidus* IPCD/DIPPS colored in teal (4NMD), Human CEPT1 colored in pink (8GYX). yCPT1 colored same as map and models shown in panel A.

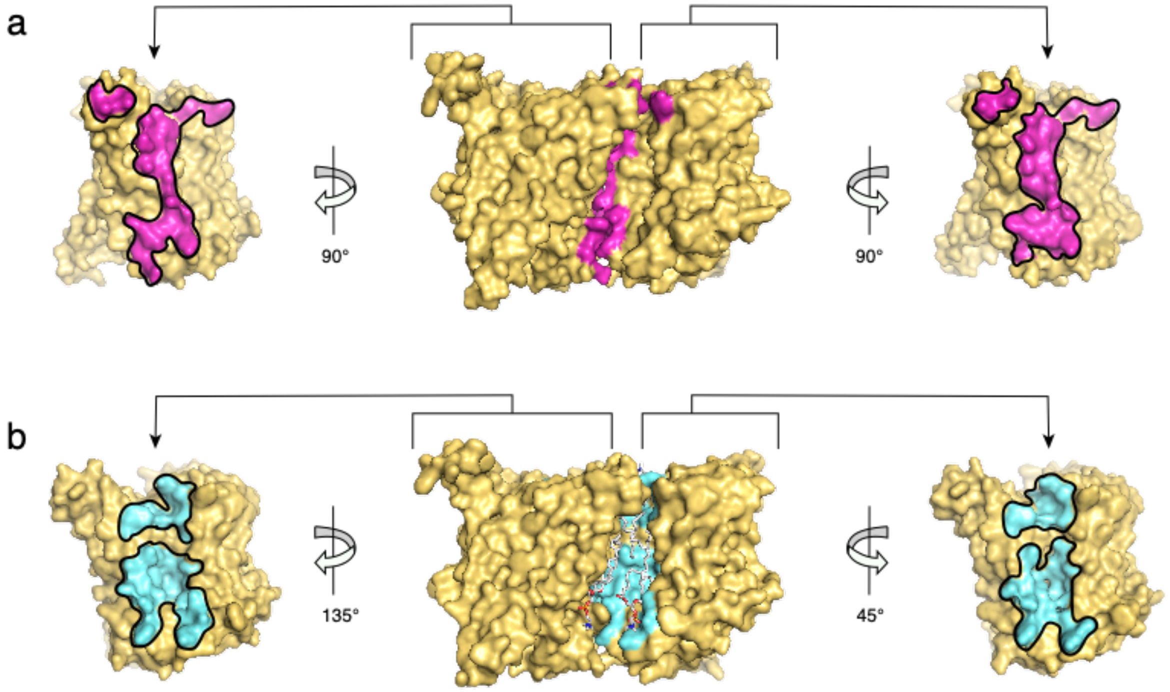

**Supplementary Figure 5. Dimerization interface of the yCPT1. a.** The dimer interface mediated through the protein interactions are shown in magenta. Interface residues are defined as the ones within 4 Å of each other. **b.** Dimer interface mediated through contact with phospholipid molecules are shown in cyan. For clarity, the lipid molecules are removed in the split subunit view. The interface residue is defined as the ones within 4Å from the neighboring subunit or the lipid molecules.

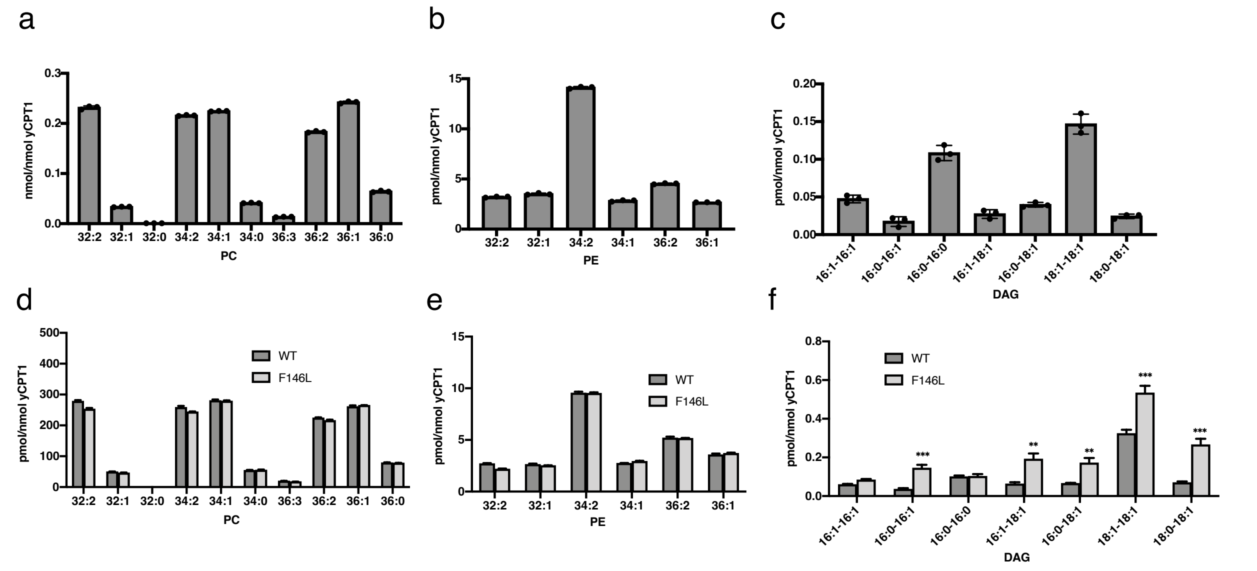

**Supplementary Figure 6. Quantification of PC, PE, and DAG from the purified proteins.** Lipids were extracted from 900 pmol of purified yCPT1 WT or yCPT1 F146L mutant. The amounts of PC (a, d), PE (b, e), and DAG (c, f) were determined by LC-MS/MS. Data are presented as the mean ± SD. ** and *** indicate $P < 0.01$ and $P < 0.001$ as compared with yCPT1 WT, respectively. Statistical significance was determined using a Student's t-test.

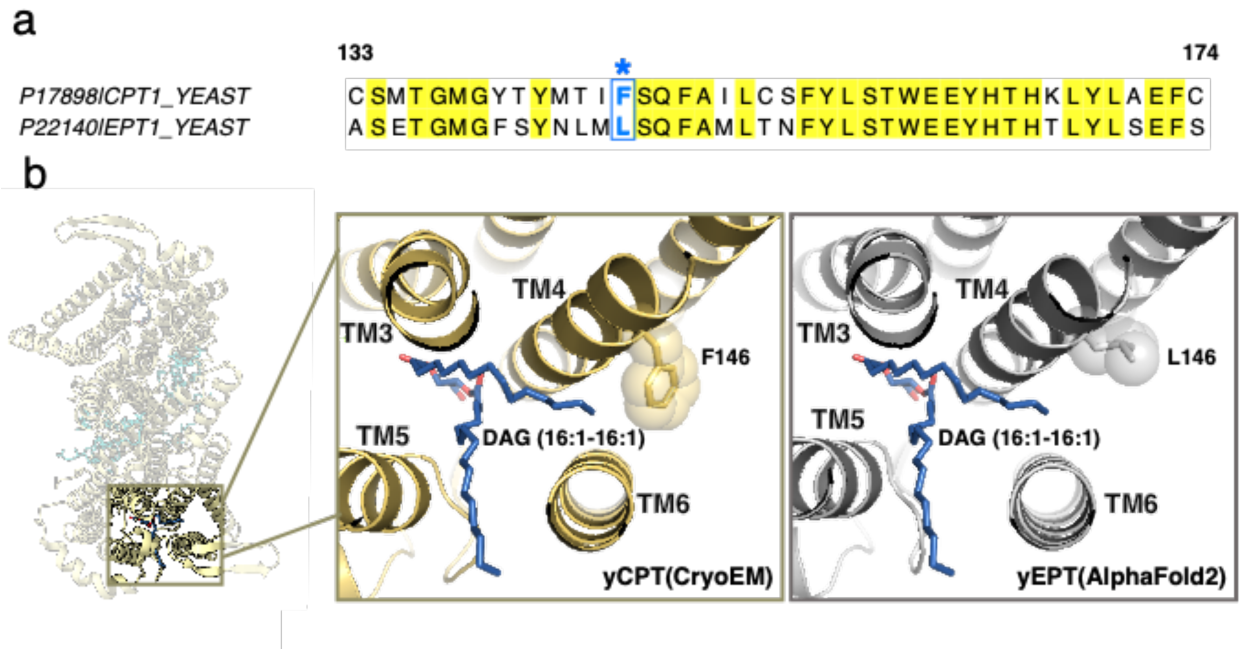

**Supplementary Figure 7. Structural comparison between the cryo-EM yCPT1 and AF2-predicted yEPT1 structures. a.** Sequence alignment of yCPT1 (UniProt ID: P17898) and yEPT1 (UniProt ID: P22140) around TM4 region. Identical residues between these proteins are highlighted yellow. yCPT1 F146 and yEPT1 L146 are marked by an asterisk and shown in blue. **b.** Comparison of the DAG-binding site of cryo-EM structure of yCPT1 and AF2-predicted structure of yEPT1.

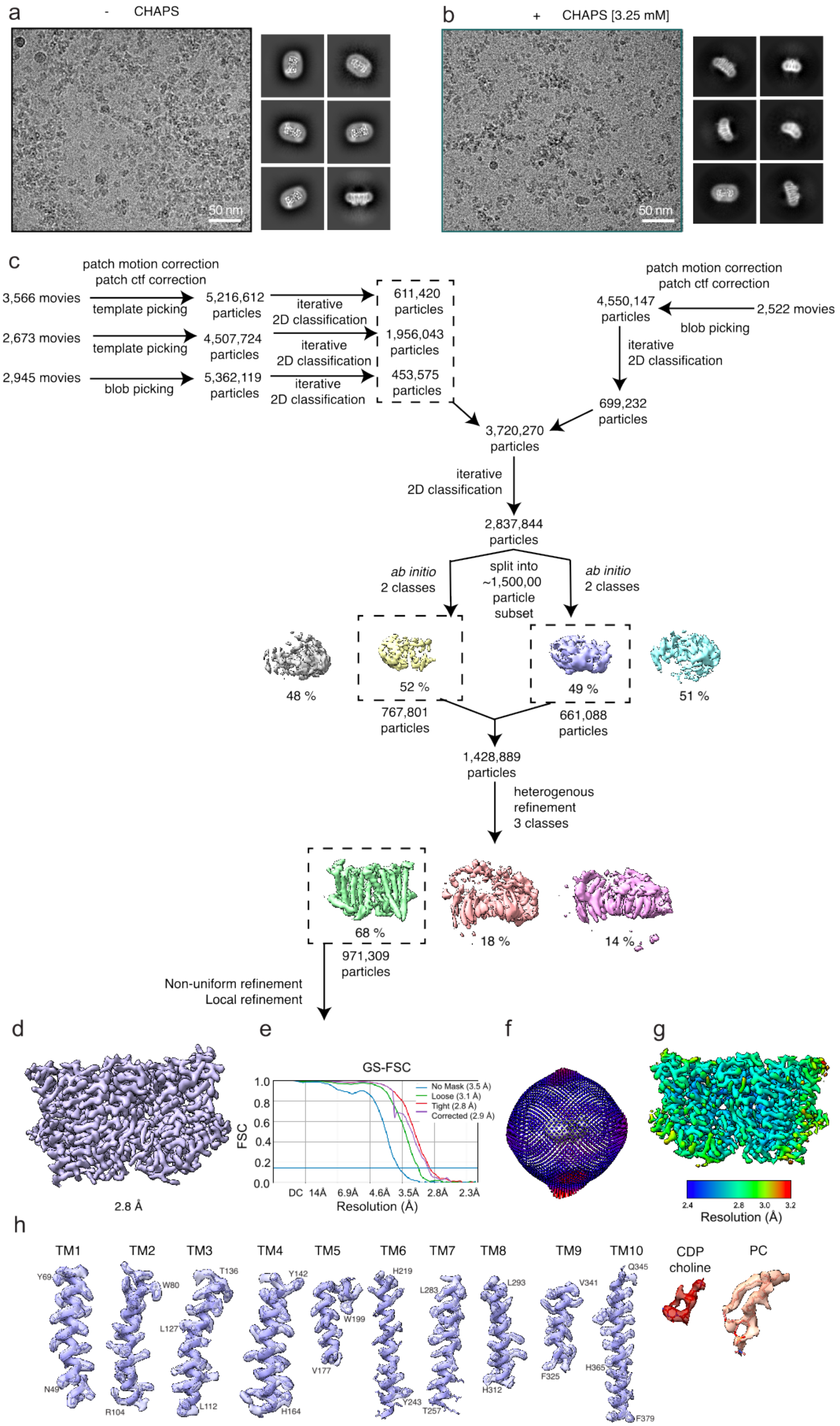

a
- CHAPS
50 nm
b
+ CHAPS [3.25 mM]
50 nm
c
patch motion correction
patch ctf correction
3,566 movies
template picking
5,216,612 particles
iterative 2D classification
611,420 particles
2,673 movies
template picking
4,507,724 particles
iterative 2D classification
1,956,043 particles
2,945 movies
blob picking
5,362,119 particles
iterative 2D classification
453,575 particles
patch motion correction
patch ctf correction
4,550,147 particles
2,522 movies
blob picking
iterative 2D classification
699,232 particles
3,720,270 particles
iterative 2D classification
2,837,844 particles
ab initio 2 classes
split into ~1,500,00 particle subset
ab initio 2 classes
48 %
52 %
49 %
51 %
767,801 particles
661,088 particles
1,428,889 particles
heterogenous refinement 3 classes
68 %
18 %
14 %
971,309 particles
Non-uniform refinement
Local refinement
d
2.8 Å
e
GS-FSC
FSC
1.0
0.8
0.6
0.4
0.2
0.0
DC 14Å 6.9Å 4.6Å 3.5Å 2.8Å 2.3Å
Resolution (Å)
No Mask (3.5 Å)
Loose (3.1 Å)
Tight (2.8 Å)
Corrected (2.9 Å)
f
g
2.4 2.8 3.0 3.2
Resolution (Å)
h
TM1 TM2 TM3 TM4 TM5 TM6 TM7 TM8 TM9 TM10
CDP choline
PC
Y69 N49 W80 R104 L127 T136 L112 Y142 H164 W199 V177 H219 Y243 L283 T257 L293 H312 V341 F325 Q345 H365 F379

**Supplementary Figure 8. Cryo-EM processing of yCPT1 in complex with CDP-choline. a. b.** Sub-area of motion corrected micrographs of vitrified yCPT1 in the presence (**a**) and absence (**b**) of 3.25 mM CHAPS. Representative class averages, box size 276 Å. **c.** Cryo-EM processing workflow. Initial particles were picked by cryoSPARC template picking, or crYOLO model as specified. 3.7 million total particles were extracted and pooled from the 4 datasets. **d. e.** After iterative 2D classification and heterogenous refinement 971,309 good particles were subjected to a non-uniform refinement reaching 2.8 Å GS-FSC resolution. **f.** Orientation distribution plot. **g.** Local resolution estimation. **h.** Density of each TM helix and CDP choline are shown contoured at 6 sigma. Density of PC is shown contoured at 3 sigma.

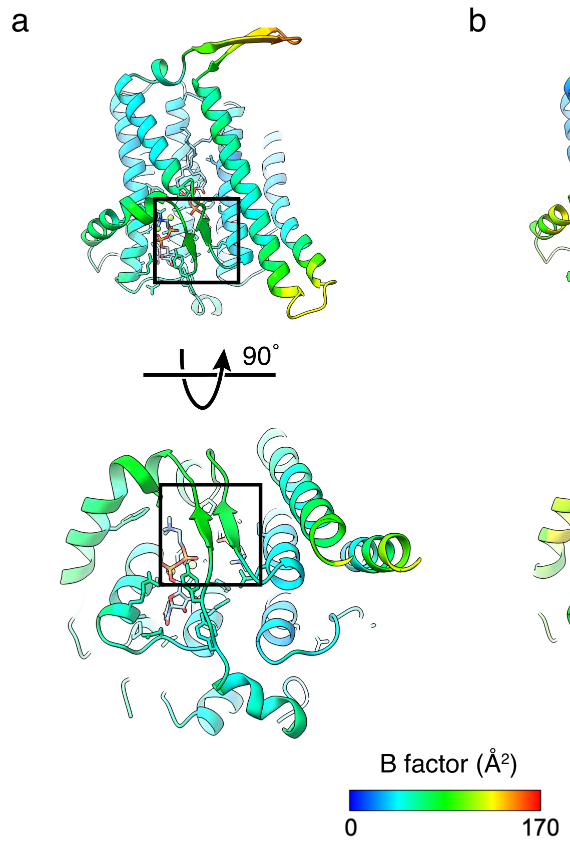

**Supplementary Figure 9. B-factor comparison between yCPT1 bound with CDP-choline/PC and DAG. a.** B-factor plot of yCPT1 bound with CDP-choline and PC. **b.** B-factor plot of yCPT1 bound with DAG alone and no CDP-choline. The enclosed area is rigidified upon binding of CDP-choline, while the same area is the most flexible region in the DAG-bound form.

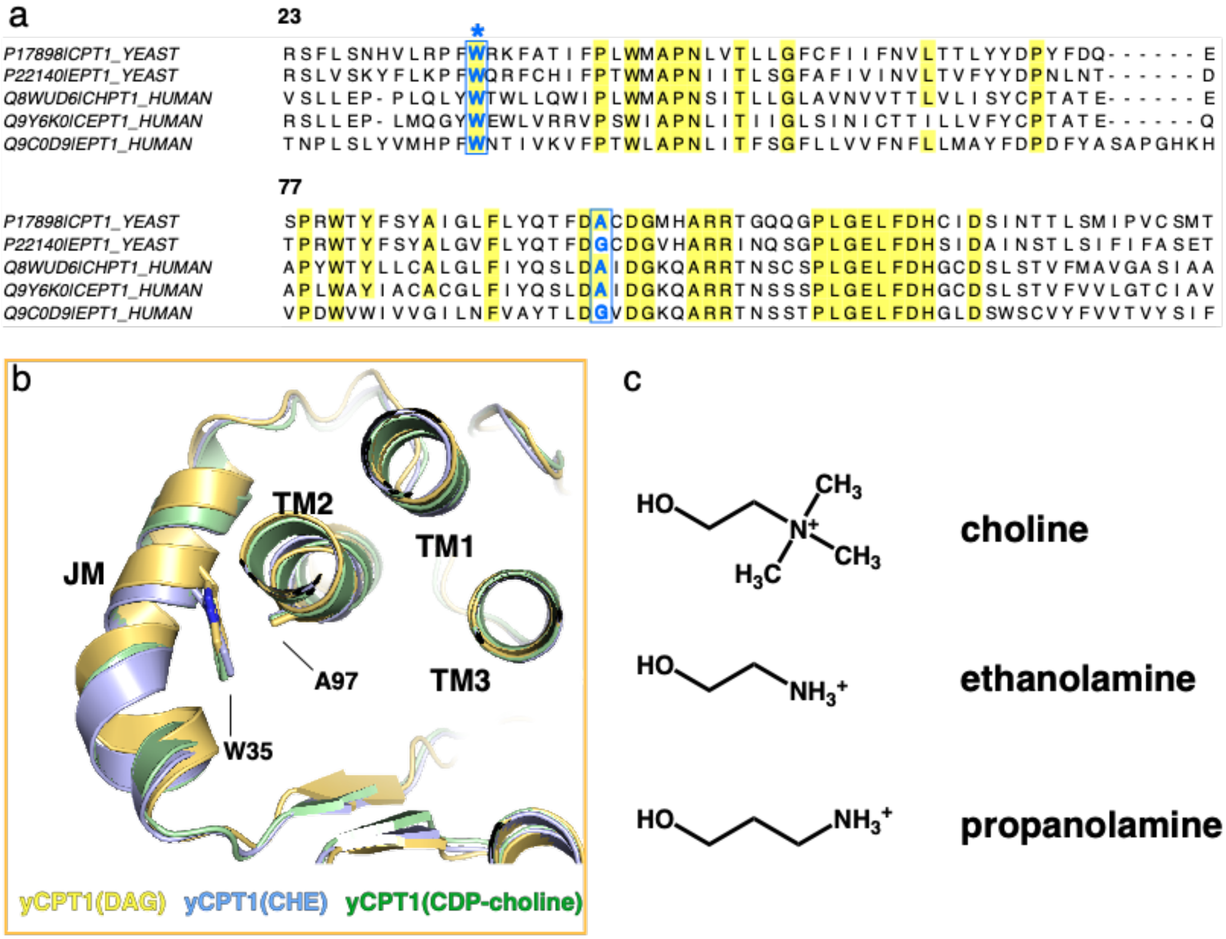

**Supplementary Figure 10. Specificity determinant for the lipid head group. a.** Sequence alignment of CDP-choline/ethanolamine transferases from yeast and human. yCPT1 (UniProt ID: P17898), yEPT1 (UniProt ID: P22140), hCHPT1 (UniProt ID: Q8WUD6), hCEPT1 (UniProt ID: Q9Y6K0), and hEPT1 (UniProt ID: Q9C0D9). Highly homologous residues are highlighted in yellow and the conserved Tryptophane in the JMH (equivalent to Trp35 in yCPT1) and the residue equivalent to Ala97 in yCPT1 are colored in blue with an asterisk marks. **b.** Structural comparison of yCPT1 in different states. Note that the position of W35 is constant throughout the states. **c.** Chemical structures of choline, ethanolamine, and propanolamine.

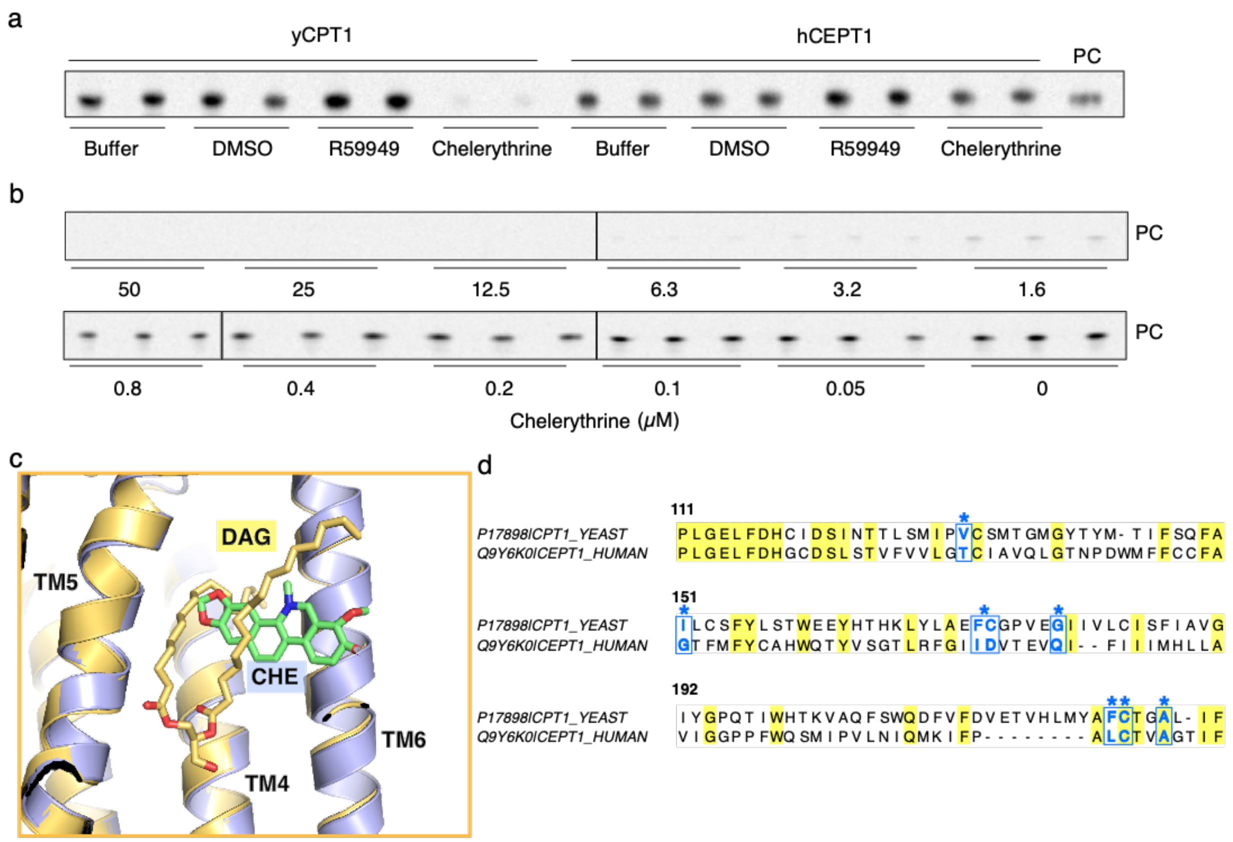

**Supplementary Figure 11. Specific inhibition of yCPT1 by chelerythrine. a.** TLC analysis of radiolabeled PC shows chelerythrine specifically inhibits yCPT1, while it doesn't suppress hCEPT1 enzymatic activity. R59949 doesn't inhibit either yCPT1 or hCEPT1. **b.** PC synthesis activity of the purified yCPT1 is suppressed by chelerythrine concentration dependently. Chelerythrine concentration is titrated from 0,05μM to 50μM. **c.** Superposition of yCPT1 structures bound to either DAG or chelerythrine. Chelerythrine occupies the DAG binding pocket and overlaps where DAG binds. **d.** Sequence alignment of yCPT1 (UniProt ID: P17898) and hCEPT1 (UniProt ID: Q9Y6K0) around TMs 4-6. Identical residues are highlighted yellow and chelerythrine-binding residues are marked by asterisk and shown in blue.

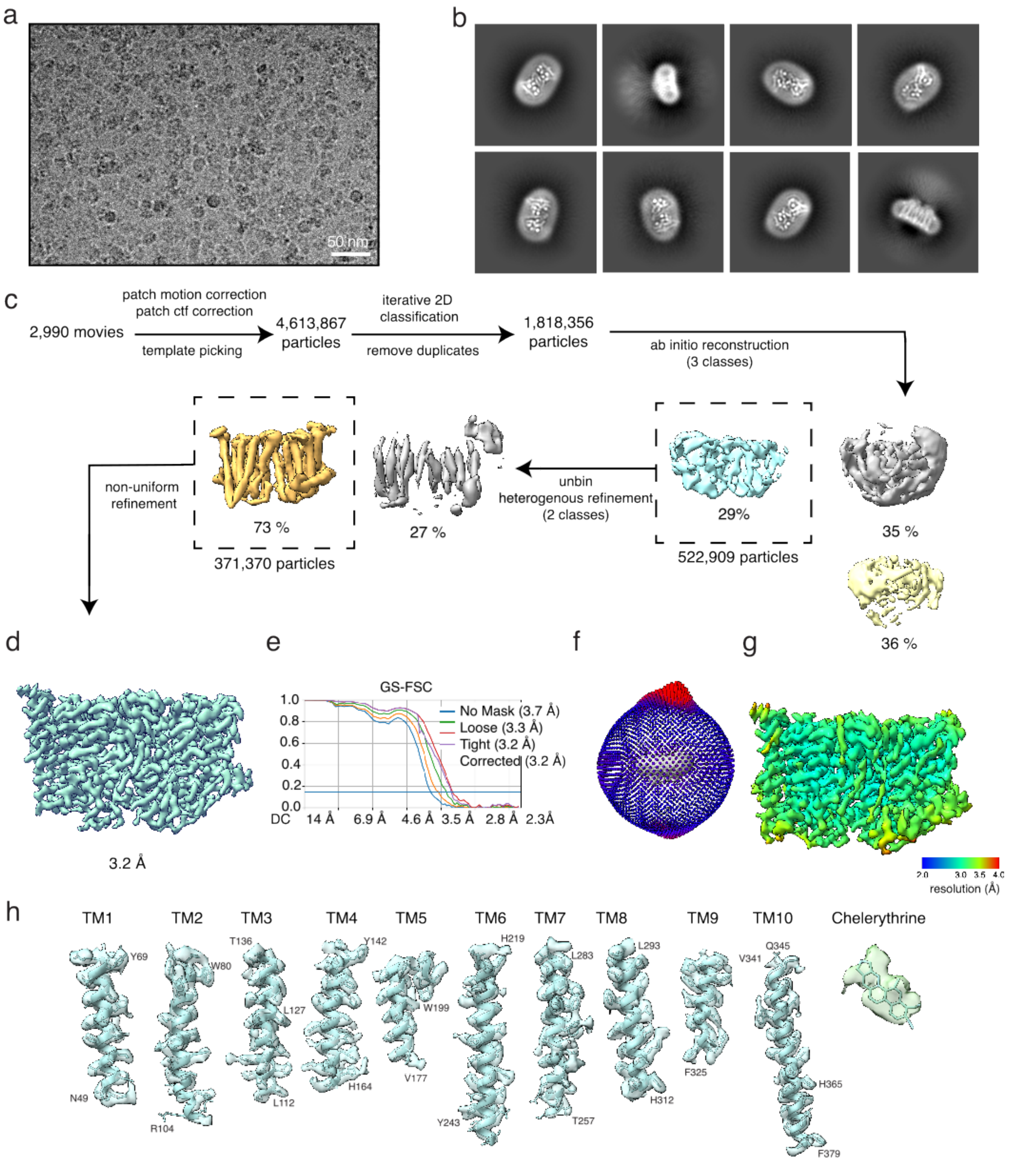

**Supplementary Figure 12. Cryo-EM processing of yCPT1 in complex with chelerythrine. a.** Sub-area of motion corrected micrograph of vitrified yCPT1 particles. **b.** Representative 2D classes. Box size 276 Å. Scale bar 50 A. **c.** Cryo-EM processing workflow. 4.6 million initial particles were picked from 2,990 micrographs using crYOLO. **d. e.** After iterative 2D classification and heterogenous refinement 371,370 good particles were subjected to a non-uniform refinement, reaching 3.2 Å GS-FSC resolution. **f.** Orientation distribution plot. **g.** Local resolution estimation. **h.** Density of each TM helix and chelerythrine are shown contoured at 3 sigma.

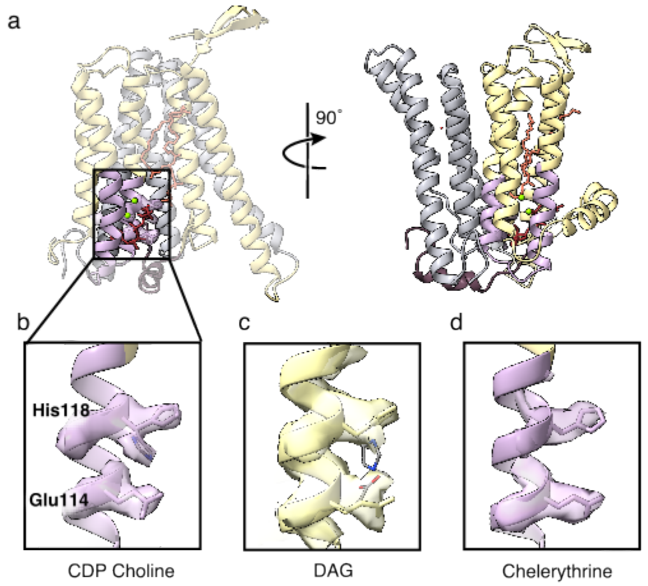

**Supplementary Figure 13. Conserved Histidine residue can adopt different rotameric conformations depending on bound substrate. a.** Overall structure of yCPT1 monomer in complex with CDP-choline. The catalytic site is highlighted in the left panel and enlarged in (b). **b, c, d.** Comparison of the cryoEM density maps and models of His118 and Glu114 of yCPT1 bound with CDP-choline (**b**), DAG (**c**), and chelerythrine (**d**).

### Supplementary Table1. Cryo-EM data collection, refinement and validation statistics

| | yCPT1 DAG (EMDB-42357) (PDB 8UL9) | yCPT1 CDP choline/PC (EMDB-42496) (PDB 8URP) | yCPT1 Chelerythrine (EMDB-42500) (PDB 8URT) |
|---|---|---|---|
| **Data collection and processing** | | | |
| Magnification | 81,000 | 81,000 | 81,000 |
| Voltage (kV) | 300 | 300 | 300 |
| Electron exposure (e–/Å$^2$) | 60 | 60 | 60 |
| Defocus range (μm) | 1.5-2.4 | 1.5-2.4 | 1.5-2.4 |
| Pixel size (Å) | 1.08 | 1.08 | 1.08 |
| Symmetry imposed | C1 | C1 | C1 |
| Initial particle images (no.) | 2,913,179 | 2,936,145 | 4,613,867 |
| Final particle images (no.) | 677,515 | 1,349,967 | 371,370 |
| Map resolution (Å) | 3.22 | 2.93 | 3.20 |
| FSC threshold | 0.143 | 0.143 | 0.143 |
| Map resolution range (Å) | 2.8-45 | 2.6-29 | 2.9-52 |
| **Refinement** | | | |
| Initial model used | AlphaFold2(P17898) | XXX | XXX |
| Model resolution (Å) | 3.20 | 2.90 | 3.20 |
| FSC threshold | 0.143 | - | - |
| Model resolution range (Å) | XXX | XXX | XXX |
| Map sharpening *B* factor (Å$^2$) | 148.9 | 130.6 | 144.4 |
| Model composition | | | |
| Non-hydrogen atoms | 6668 | 6747 | 6622 |
| Protein residues | 774 | 774 | 774 |
| Ligands | 14 | 16 | 14 |
| *B* factors (Å$^2$) | | | |
| Protein | 62.50 | 62.61 | 76.47 |
| Ligand | 62.82 | 72.25 | 77.88 |
| R.m.s. deviations | | | |
| Bond lengths (Å) | 0.003 | 0.002 | 0.008 |
| Bond angles (°) | 0.623 | 0.455 | 1.017 |
| Validation | | | |
| MolProbity score | | | |
| Clashscore | 10.24 | 5.50 | 6.30 |
| Poor rotamers (%) | 0.15 | 1.60 | 0 |
| Ramachandran plot | | | |
| Favored (%) | 94.55 | 97.14 | 97.01 |
| Allowed (%) | 5.45 | 2.86 | 2.99 |
| Disallowed (%) | 0 | 0 | 0 |
